# Keeping up with the genomes: efficient learning of our increasing knowledge of the tree of life

**DOI:** 10.1101/758755

**Authors:** Zhengqiao Zhao, Alexandru Cristian, Gail Rosen

**Affiliations:** Department of Electrical and Computer Engineering, Drexel University, Philadelphia, PA, U.S.; Department of Computer Science, Drexel University, Philadelphia, PA, U.S.

## Abstract

It is a computational challenge for current metagenomic classifiers to keep up with the pace of training data generated from genome sequencing projects, such as the exponentially-growing NCBI RefSeq bacterial genome database. When new reference sequences are added to training data, statically trained classifiers must be rerun on all data, resulting in a highly inefficient process. The rich literature of “incremental learning” addresses the need to update an existing classifier to accommodate new data without sacrificing much accuracy compared to retraining the classifier with all data. We demonstrate how classification improves over time by incrementally training a classifier on progressive RefSeq snapshots and testing it on: (a) all known current genomes (as a ground truth set) and (b) a real experimental metagenomic gut sample. We demonstrate that as a classifier model’s knowledge of genomes grows, classification accuracy increases. The proof-of-concept naïve Bayes implementation, when updated yearly, now runs in 1/4^*th*^ of the non-incremental time with no accuracy loss. In conclusion, it is evident that classification improves by having the most current knowledge at its disposal. Therefore, it is of utmost importance to make classifiers computationally tractable to keep up with the data deluge.

## Introduction

Recent advances in genomics have resulted in exponential increases in the rate at which data are collected. Inspired by [1], we visualize the growth of National Center for Biotechnology Information (NCBI) Reference Sequence (RefSeq) Bacterial genome database [2, 3] in Figure 1. Figure 1A shows the total number of complete genomes added/updated in the database in yearly increments until March 2nd 2019^1^, and Figure 1B shows the number of new and updated completed genomes every year. As shown in Figure 1, there are now thousands of genomes being sequenced per year, providing vital information for understanding prokaryotic species diversity, with major efforts like the Genome Encylopedia of Bacteria and Archaea contributing to this expansion [4].

**Figure 1.**
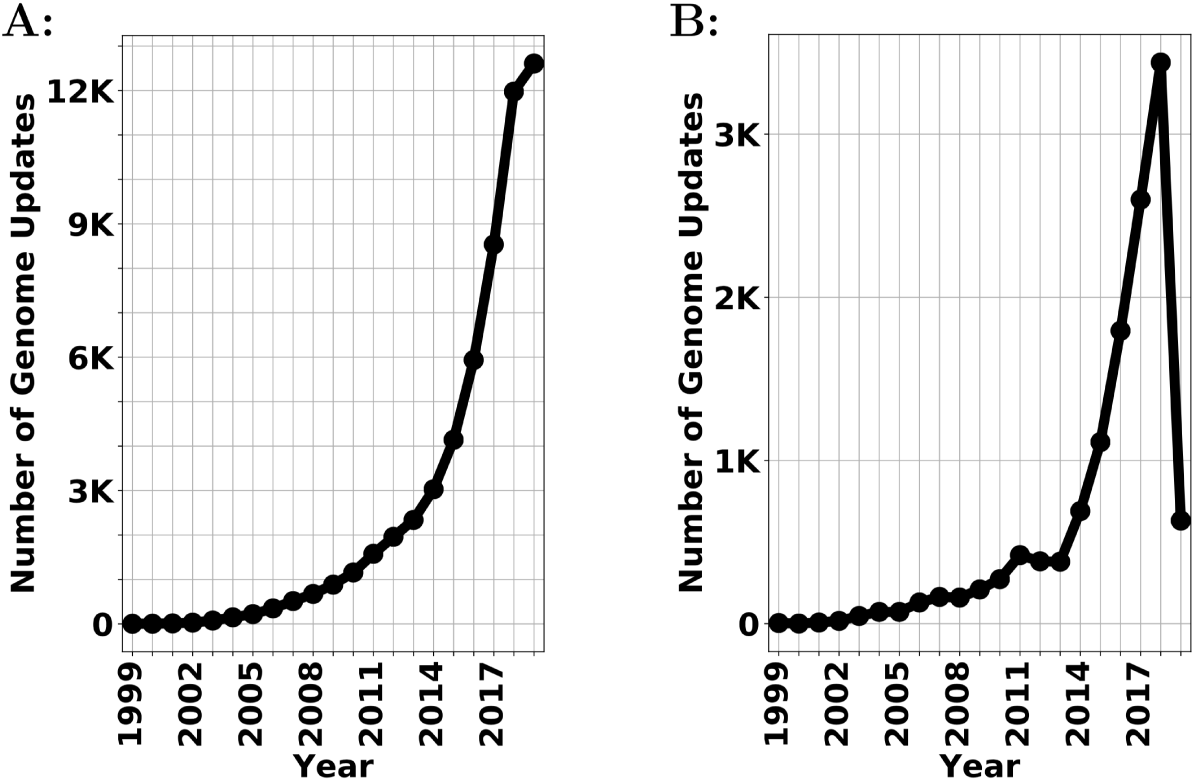
Number of updates in the NCBI Bacteria Genome database. **A:** Accumulative number of genomes updates per year; B: compared with last year, the number of new updates per year

Moreover, the cost of sequencing DNA is reducing at a rate that is outpacing Moore’s law [5]. In other words, the increase of computational power cannot keep up with the growth of the number of genomes (as well as metagenomes) being added.

One of the main challenges in metagenomics is to answer, “Who is there?”: identifying the microorganisms in the sample. Taxonomic classification – classifying the reads in metagenome sequencing data – uses aligners, read mappers, classifiers and other “base techniques” to solve this problem [6–10]. Taxonomic classification is usually one of the first steps in a metagenomic pipeline [11]. Once these organisms are identified, they are then used in downstream analyses, such as alpha/beta diversity measures, ordination, feature selection, phenotype classification, etc.

However, while many methods have been proposed for taxonomic classification [12, 13], the accuracy of these methods using different training databases has not been fully tested. This is an important issue. Because as new genome data are generated, training data sets, such as the commonly used the NCBI Reference Sequence Database (RefSeq), will change over time. Nasko et al. [14] recently demonstrated that more reads are classified (as opposed to be assigned to an unclassified/unknown class) by the Kraken classifier with newer database versions. Nasko’s analysis also suggested that changes in RefSeq over time may influence but not necessarily reduce the misclassification rate, with genus/species false positives shown to be 1% and 8% respectively. These metrics were, however, calculated only on a selection of 10 genomes. Accordingly, given that there is an ongoing rapid expansion of genomic data of microbial diversity, it is imperative to update taxonomic classifiers as new genomes/genes are discovered. Simply failing to update the model will result in lower accuracy due to incomplete knowledge. In addition, the way that most researchers train their taxonomic classifiers is a static process: A fixed number of genomes are used as training data, and when more reference genomes are added to NCBI, the classifiers must be completely retrained to accommodate the new data. This type of training is computationally unsustainable, because the cost of sequencing DNA is reducing at a rate that outpacing Moore’s law [5].

The traditional model of retraining of classifiers, each time training on the entire dataset, cannot sufficiently keep pace with new discoveries. One possible course of action would be to update the database less frequently. However, the resulting analysis will not be as “accurate” as it could be if frequently updated with knowledge about new genomes. As a result, there is in a need for new innovations in updating classifiers.

In RefSeq database, we can observe a new genome being added, removed and updated and its taxonomic labels can be merged with, changed to other labels and even removed from the database. In our work, we only focus on the case that new genome are added to the database. However, our algorithm can also be extend to handle the taxonomic nodes merging scenarios (see our discuss in Incremental Naïve Bayes Classifier).

We aim to solve how we can update a taxonomic classifier quickly with only new data without reprocessing the whole database. An intuitive solution is to update the training model for new data only, instead of reprocessing the whole database. In this way, we can greatly reduce the amount of redundant computations to compute more frequent updates. The naïve Bayes classifier is a natural solution, since only the columns (of the *k*-mer frequency table) of the classes being updated or added need to be changed. We demonstrate that the incremental learning algorithm allows computers to “continually learn” new information seamlessly and alleviate the current inefficient and inadequate retraining practices. We also show that the limitation of navïve Bayer classifier: its classification results can be biased by species classes that have many genomic examples (i.e: well-represented species). In this paper, we show that incremental learning algorithm is a promising solution. We developed a proof-of-concept incremental implementation of a naïve Bayes classifier. We then show that the taxonomic classification task can benefit from learning incrementally from the NCBI dataset in respect to two aspects:

- the classifier gains accuracy over the time by constantly updating the classifier with the latest knowledge (genomes).
- the time of update can be greatly reduced by incremental learning.

While some of the results show that NBC might not the best choice for taxonomic classifier, we hypothesize that other algorithms may be afflicted by this class bias as well. So, our work serves as a lesson.

Therefore, the main aim of this paper is to show how incremental learning can reduce computational time and enables constant improvements in accuracy through frequent updates over time. The paper highlights the importance of incremental learning and its promise, rather than the NBC classifier selection itself. Much future work remains to examine how other taxonomic classifiers can be “incrementalized”.

In this paper, “incrementalized” version of naïve Bayes taxonomic classifier demonstrates how the computational time of training is reduced while still retaining the same accuracy as the original implementation [6]. To be specific, when a new labeled sequence is available, we update the model to learn new information: If the new sequence is from an existing class, then the class will be updated based on Bayes rule; otherwise, a new class is created to accommodate this new organism. Therefore, we do not have to create a new model from scratch and process the whole database. Instead, we only need to process the new data to update the model. In addition, we present an implementation of NBC that optimizes RAM and the number of cores via a smart loading scheme that is scalable for various architectures. Our implementation of NBC is open source here: https://github.com/EESI/Naive_Bayes. Our contribution in this paper is three-fold:

- show quantitative classification results that improve over time by training a classifier on progressive RefSeq snapshots and testing the classifier on simulated reads using 5-fold cross-validation.
- qualitatively demonstrate how the classification composition of a real metagenomic gut sample changes over time by training a classifier on RefSeq dataset in 5 different years.
- propose a proof-of-concept incremental implementation of Naïve Bayes taxonomic classifier (NBC++), that can be efficiently updated with new information without having to reprocess the existing database, which drastically reduces the training time of the classifier when new data is available compared to the training time of the non-incremental implementation.

## Related Work

Our work relates to both incremental learning and microbial taxonomic classification fields. We develop and evaluate an “incrementalized” version of the naïve Bayes taxonomic classifier [6].

### Related Work on Incremental Learning

In an incremental learning setting defined by [15], a dataset 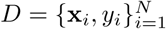 is divided into *T* parts *D* = {*D*_1_, …, *D*_*T*_}, which arrive sequentially [15]. The goal is to build an efficient algorithm that takes as input (a) a model, **M**_*t*_ that has been trained on the first *t* units of data {*D*_1_, …, *D*_*t*_} and (b) new data, *D*_*t*+1_, to then output an updated model **M**_*t*+1_. The efficiency of the algorithm can be achieved without direct access to the previous training data, {*D*_1_, …, *D*_*t*_}, while performance is maintained without catastrophically forgetting the previous model [16]. Ensemble learning algorithms can be used to achieve incremental learning. [17] proposed an incremental learning algorithm, Learn++, that ensembles “weak” classifiers during the training. New classifiers will be added to learn and explain the new and misclassified examples. Support Vector Machine (SVM) are “incrementalized” by many researchers [18–20]. SVM finds support vectors and determines a decision boundary that best separates the support vectors. A set of support vectors can be viewed as a good abstraction of training data, therefore, the existing training database can be compressed using support vectors. When the data is presented to the learning algorithm sequentially in batches, one can compress the data of the previous batches to their support vectors. Then, for each new batch of data, a SVM is trained on the new data and the support vectors from the previous learning step [18]. Deep learning approaches have been producing state-of-the-art results yet they continue to suffer from catastrophic forgetting, a dramatic decrease in overall performance when training with new data added incrementally [16]. [15] utilize Bayesian approach to update the deep learning model with new data. [16] uses new data and a small exemplar set from the old classes to update the model. Both methods show improvement on performance for incremental deep learning models. [15] works for models that have a determined number of classes which remains the same over time whereas [16] handles the cases that novel classes can be available when training. Typically, there are two types of incremental learning situations: 1) new training data *D*_*t*+1_ contains data from existing classes only; 2) new training data *D*_*t*+1_ has samples from both novel classes (new classes that were not known to the existing model) and existing classes [21]. In our application, since the amount of novel bacteria are continually and increasingly being sequenced (see the supplementary file S4_NEW_UNIQUE_SPECIES_UPDATES_NUM.pdf and other S4_NEW_UNIQUE files in “S4Figure” folder), our incremental learning algorithm should must be able to handle the increasing amount of novel classes being added to the database.

### Related Work on Taxonomic Classification

16S ribosomal RNA is useful for placing organisms on the phylogenetic tree [22], and the widely-used Ribosomal Database project classifier [23] uses a naïve Bayes classifier to provide taxonomic assignments for microbial 16S rRNA sequences. However, 16S rRNA amplification is often insufficient for discrimination at the species and strain levels of classification [12] because 16S rRNA genes are too slowly evolving to reliably separate the validly named species [24, 25]. Whole-genome shotgun sequencing provides researchers’ access to more genomic content of organisms and thus, can yield finer taxonomic resolution. Therefore, in this paper, we focus on metagenomic taxonomic classification that classifies metagenomic reads to species level taxa [6–8, 26]. Basic local alignment search tool (Blast) [26] is one of best methods for assigning a taxonomic label to an unknown sequence [7, 12]. Other Blast-based algorithms further improve the taxonomic classification accuracy by incorporating other information (e.g. last/lowest common ancestor) [27]. In incremental learning scenario, when new genomes are added to the reference database, blast database should be updated to accommodate the new genomes. Currently, Blast databases are daily updated^2^. However, Blast is not designed for high-throughput metagenomic reads classification, and its computationally expensive to get local alignments for hundreds of thousands and millions of reads. In addition to alignment based tools, there are some successful k-mer composition based tools. LMAT [8] provides a *k*-mer based scalable metagenomic classification by creating a taxonomy/genome index. Kraken [7] uses exact-match of *k*-mers, rather than inexact alignment of sequences to perform taxonomic classification. The classification is based on a well organized *k*-mer to lowest common ancestor (LCA) database and it is very efficient to search k-mers in the database. The authors show that Kraken’s accuracy is comparable to Blast based sequence classifiers and and run faster than those competing programs. When there are new genomes added to the database, Kraken needs to be trained from scratch. Although Kraken is not designed for quick updates, it is potentially “incrementalizable”: only a fraction of the taxonomy tree (the nodes of updated/newly added genomes and their ancestral nodes) and the Kraken k-mer database need be updated. NBC [6] is a machine learning based approach for taxonomic classification. It uses Naïve Bayes classifier (NBC) to classify all metagenomic reads to their best taxonomic match and is good at estimating the number of reads of particular organisms that are known to the classifier shown via *l*1-norm distance and log-modulus deviation [12]. The NBC classifier is, by nature, “incrementalizable”. When new genomes are added, the classifier can be updated with only the conditional probabilities of the new species. In recent years, there are novel tools that incorporate deep learning techniques to perform taxonomic classification [28–30]. However, the deep learning models can’t be quickly updated in case of new species. The final dense layer has to be modified to accommodate the new species. The update speed can be improved by training the new model with weight from the existing deep learning model. In this paper, we “incrementalize” the NBC classifier to show that we can use less computation while retaining the same accuracy and demonstrate the efficacy of our implementation over increasing knowledge.

## Results and Discussion

In this section, we use NCBI Reference Sequence (RefSeq) Bacterial genome database (see S4 in Supporting information section for details) to evaluate: 1) the k-mer size influence on taxonomic classification accuracy; 2) the incremental taxonomic classification performance; 3) the dynamics of a classifier trained on different yearly snapshots of the RefSeq bacterial database and its influence on taxonomic profiling results. These results demonstrate the necessity of incremental learning for metagenomic taxonomic classification.

It is important to note that unlike the original implementation of NBC [6], where each read was first classified on the genome-level, each read here is classified to a species class. Each species’ k-mers are aggregated and trained to be a “species class”.

In the following results, we show that an incremental learning classifier has the following advantages over the traditional non-incremental version:

- incremental learning can help when a computer has limited memory and cannot process an entire large training dataset at-once – by incrementally processing small batches of the training data at a time.
- the classifier can keep up with the latest knowledge (and associated accuracy gains) efficiently without the cost of a full retrain.

### Evaluating NBC on a larger dataset: *k*-mer size vs. accuracy

Benefiting from the incremental learning ability, the entire training dataset does not need to be loaded into memory. Instead, we can simply load training data in small batches and continuously update our classifier. Therefore, NBC++ can easily use large sizes of *k*-mers. In this experiment, we use 5-fold cross-validation to evaluate the effects of *k*-mer size on taxonomic classification accuracy. Figure 2 shows the average species level accuracy as a function of k-mer size (the standard deviation is not signification thus removed from the visualization). Because we split our training and testing data on genome level (i.e., we evaluate our classifier on genomes/strains that the classifier have not seen in the training phase. The labels of those genomes/strains may or may not have the same species level label as those in training data), there is a chance that the testing reads are simulated from genomes that are unknown to the classifier. The dotted line shows the accuracy on all testing reads (including reads that are unknown to the classifier) and the solid line shows the accuracy on only the known testing reads. The limitation of NBC is shown in Figure 2, where the accuracy does not improve past mers. Please refer to section “Mistakes: Current implementation’s tendency to classify to well-represented species classes” for more detailed discussion about misclassification found in the NBC implementation.

**Figure 2.**
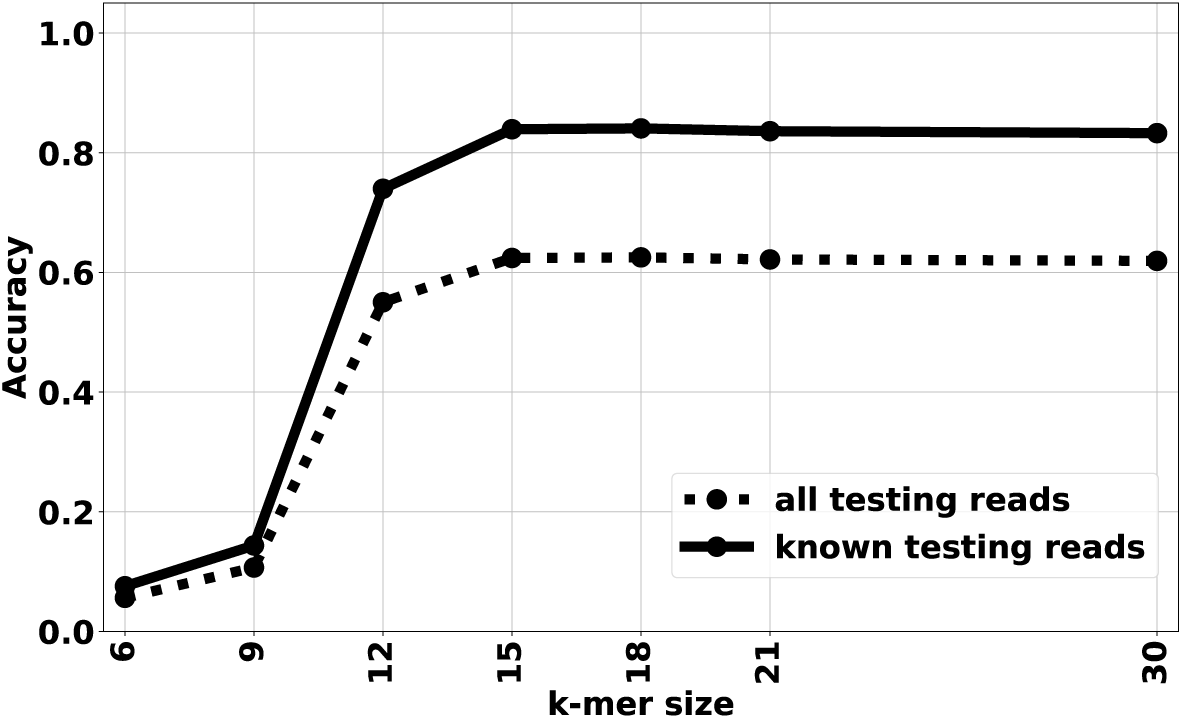
The species-level NBC++ accuracy vs. k-mer size.

To obtain upper levels of taxonomic classification, the species-level of classification is “traced back” from the species label using the NCBI taxonomic database provided by [2]. Therefore, we do not do any class training at these levels. All classifications are done at the *species level*. Figure 3 shows the average accuracy on 6 taxonomic levels: species, genus, family, order, class and phylum, as a function of k-mer size evaluated on known testing reads. Note that species level accuracy is around 84% and the genus level accuracy is around 92%. As the taxonomy level of classification increases, the accuracy increase accordingly. This indicates that although NBC++ can misclassify some testing reads to a wrong species, it may be correct on genus/class/phylum/etc level.

**Figure 3.**
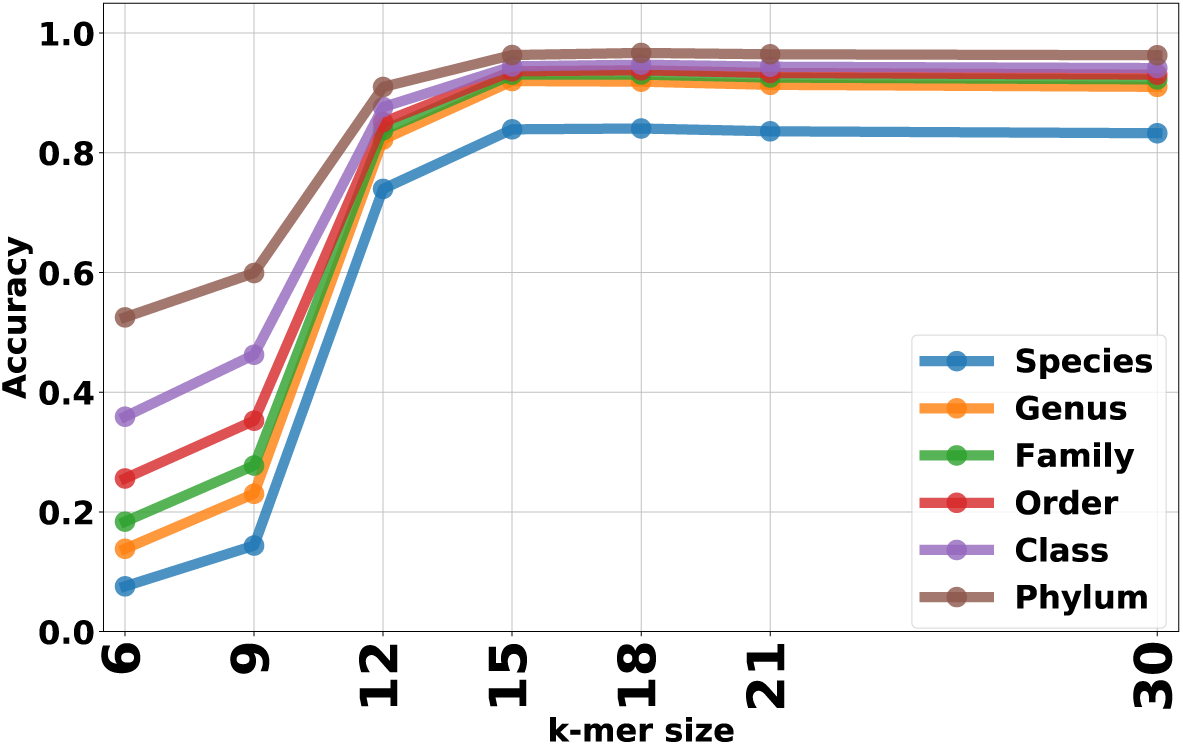
NBC++ accuracy on different taxonomic levels vs. k-mer size. Each read is classified on the species level, and higher-level taxonomic levels are calculated by “trace back” through the NCBI phylogenetic tree

### NBC++ taxonomic classification performance

To evaluate NBC++ while also demonstrating it on a real dataset, we designed experiments to simulate sequences being added yearly to the NCBI Bacterial database. In Figure 4, the species-level classification accuracy is calculated for reads simulated from the early-2019 database, with new genomes being sequenced and adding/updating species to the training database each year. From Figure 4, we can see a trade-off of more knowledge improving classification (because the class is now known) as opposed to a classifier getting “confused” between classes (as more and perhaps similar classes are added). As more data are being added, the known-species classification accuracy goes down but reaches a stable level (yellow curve), but this loss in “known” performance is dwarfed by the fact that overall accuracy (blue curve) is a function of classes that are “known” (green curve). More specifically, in 2002, as more genomes are added, NBC++ begins to misclassify some species since more are in the database, resulting in a drop to a little over 80% accuracy by 2005; however, the species that it is classified to has near 90% accuracy of being from the correct genera (see Fig. 5). This loss in performance is overshadowed when we look at the effect of the database version when using all current taxa. The overall accuracy (blue curve) is a function of the amount of taxa known at that year, and as we know more, the accuracy increases. In fact, due to our 5-fold cross-validation, there are many species with less than 5 examples of a species, and therefore, it is marked as unknown if it is not in the training set, resulting in a little over 60% accuracy by 2018. This shows that even if a species is known, it still doesn’t have enough examples to fully train the diversity of that species. Overall, the incremental implementation can gain the valuable information added per year.

**Figure 4.**
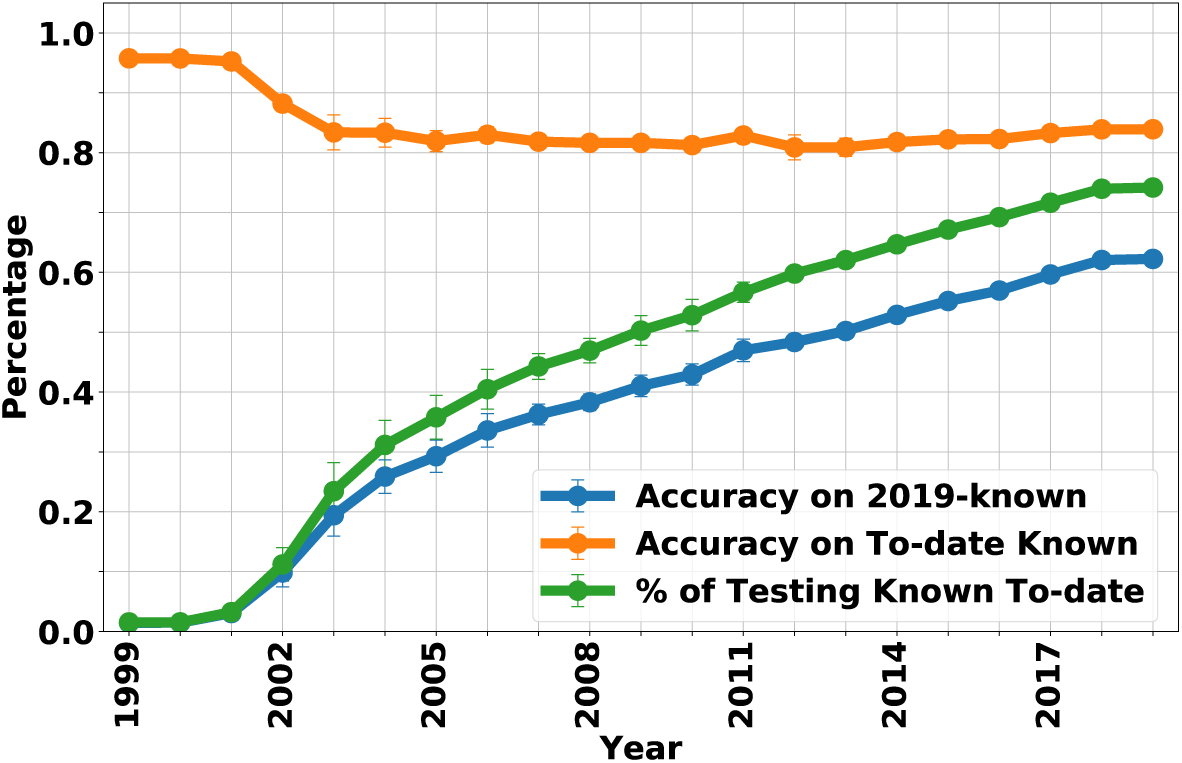
NBC++ species-level accuracy is evaluated on simulated reads reads from different species. The x-axis represents each year, where the training database is only the genomes added/updated that year. We show that the accuracy on reads from known species up-to-that-year has more than 80% accuracy and has a drop as more closely related species are added (yellow curve). The accuracy on all 2019-known species (blue curve) increases as a function of the percentage of reads from to-year known species (green curve). Therefore, the accuracy is only as good as the knowledge in our database. If for example, we only have knowledge of 3 species (which was the case in 1999), the accuracy is poor since it is testing on the thousands of species known today (see the S4_CUMULATIVE_SPECIES_UPDATES_NUM.pdf in “S4Figure” folder.)

**Figure 5.**
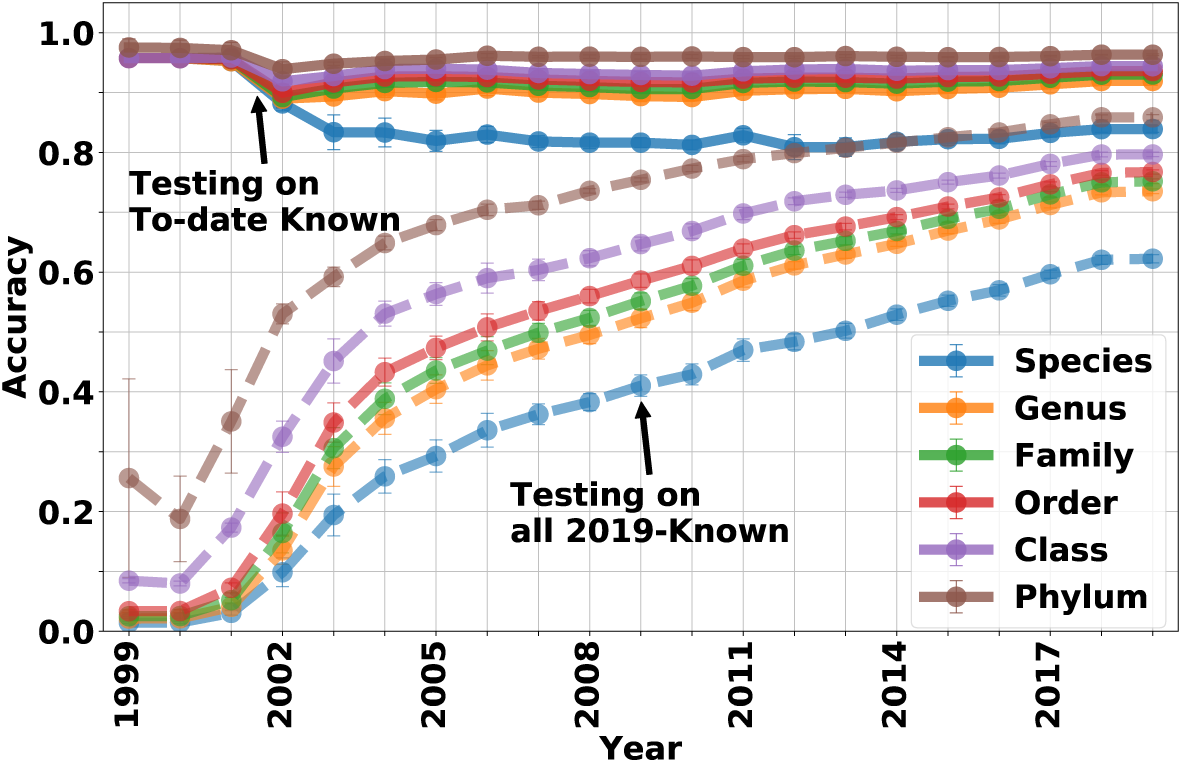
NBC++ accuracy on all taxonomic levels is evaluated on simulated reads reads from different species. The accuracy of NBC (assigned on the species-level) on genomes known to-the-year are shown in solid lines and for the entire current set of testing reads is shown in dashed lines. As more genomes are added, there is ambiguity confusing the classifier (most notably on the species level), but this performance loss is small compared to the fact that as more data is known, the better the performance.

Figure 5 shows the accuracy of NBC++ (assigned on the species-level) for genomes known to the year that is classified (solid) and on all 2019-known genomes (dotted) on 6 taxonomic levels (predictions on taxonomic levels higher than species are inferred from the species level predictions). From the figure we can see that as more genomes are added, there is ambiguity confusing the classifier but this is small compared to the fact that as more data is known, the better the classification become.

Figure 6 shows the training time of NBC++ on the database. We compare the NBC++ incremental learning feature with the non-incremental learning setting (shown as NBC in Figure 6). Although the full training time of traditional NBC is over 16 hours with 15 threads for data up to 2018, the incremental version using all new 2018 genomes to update the existing model from 2017 would only take ∼5 hours with 15 threads. In 2019, we only have new genomes until March 2nd. Therefore, the update time is significantly smaller for NBC++ whereas NBC has to reprocess the whole database again and results in a waste of computation time.

**Figure 6.**
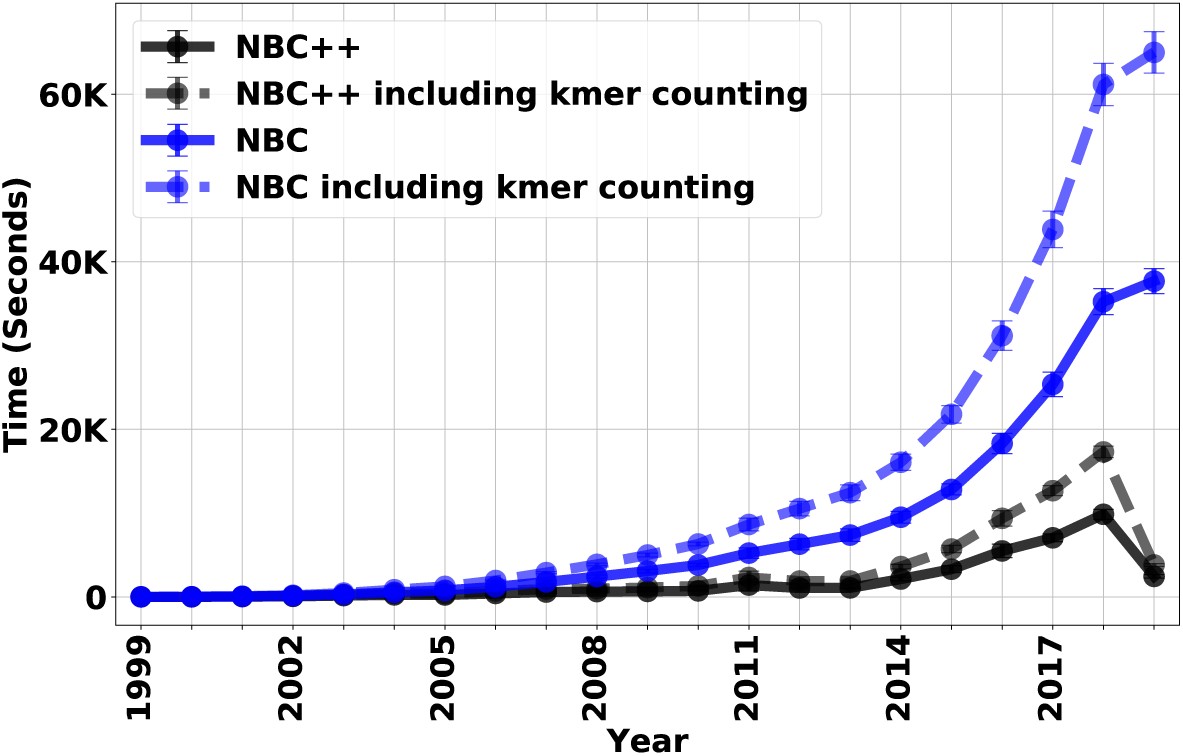
NBC Incremental learning classifier (NBC++) training/updating time is evaluated by the same simulated datasets. This figure shows the update time for incremental learning method (in blue) and traditional full update (NBC) method (in black). The different components of the algorithm are shown – the time that it takes for the core NBC algorithm (solid) vs. that time plus the average time per genome for Jellyfish to count k-mers (dashed). We can see that the k-mer counting alone can start to add a significant amount of computation when recomputing the entire training dataset.

Now we discuss the cost of model updates in a broader practical scenario. We are given two options to keep our model up-to-date. The first option is to train a model from scratch whenever an updated model is needed. This option requires to maintain a database because training a model from scratch requires both old and new data. And the second option is to update the model with new data only. This option requires the model to be stored in the system for future updates. However, in the best case scenario for the first option, we don’t need the access to the model between updates, therefore we don’t have to pay for the storage of the model for option 1. Since we need to pay for the raw data storage in option 1 and for the model storage in option 2, we can assume that the storage cost of two options are the same (usually it is not the case because the model size is often smaller than the raw data because model can be considered as a form of abstraction of the raw data). Then the only cost is the computational cost for model update. According to Amazon Web Services (AWS), a “m5.8xlarge” EC2 computing node with 32 cores and 128GB memory costs 1.536 dollars per hour^3^. If it takes 16 hours and 5 hours for option 1 and 2 to run respectively. Option 2 can save 16.896 dollars for only 1 update. If the model is updated daily, at least 6167.04 dollars can be saved without considering the increase of training time when more and more data are added to the database.

### Evaluation of NBC++ on a healthy gut sample taxonomic profiling results over time

Nasko et al. [14] uses various experiments to show that Kraken (and derivatives like Bracken) are able to classify more organisms as the RefSeq database grows. Here, we show that NBC++’s classification results change as the database size grows. Note that while NBC++ does not mark anything as “unclassified”, its classifications should get more accurate. To demonstrate how up-to-date knowledge affects results, we design real NCBI-over-time experiments to show how the results of NBC++ can be influenced by the training data.

We demonstrate using a time-progressing NCBI training database for NBC to classify a fixed human fecal (a.k.a. gut) sample (SRA ID: SRS105153), which is the same sample used in Figure 5b in [14]. For comparison, the NCBI taxonomic classification of this sample can be found online at: https://trace.ncbi.nlm.nih.gov/Traces/sra/?run=SRR514214&krona=on&dataset=0&node=0&collapse=true&color=false&depth=10&font=11&key=true. We also train 5 models using all RefSeq bacterial genomes in NCBI. In Nasko et al., 9 versions of the RefSeq database are used, but the exact results are not discussed – only the level of taxonomy that could be resolved and how many reads remained unclassified. In our experiments, we use the NCBI RefSeq bacterial genomes using 5 database divisions and show how the classifications change over time (with NBC++ labeling everything). The first division is composed of genomes added during 1999, the 2nd division are genomes added during 2000 to 2004 (version 2004), the 3rd division are genomes added during 2005 to 2009 (version 2009), the 4th division are genomes added during 2010 to 2014 (version 2014), and the the 5th division are genomes added during 2015 to 2019 (version 2019), respectively. The composition change on different taxonomic levels are evaluated for each of these divisions. Figures 7 and 8 show the predicted bacteria relative abundance of the gut sample over time on the genus and phylum levels, respectively (with species and order levels in the appendix). Taxa with lower than 5% relative abundance are moved to either “Others: Old” (dark blue, top of the bar) or “Others: New” category (light blue, bottom of the bar) category depending on whether the taxa are recently added or were added in previous years. The training database version that contains the taxa shown in the bar plots are labeled on the bars.

**Figure 7.**
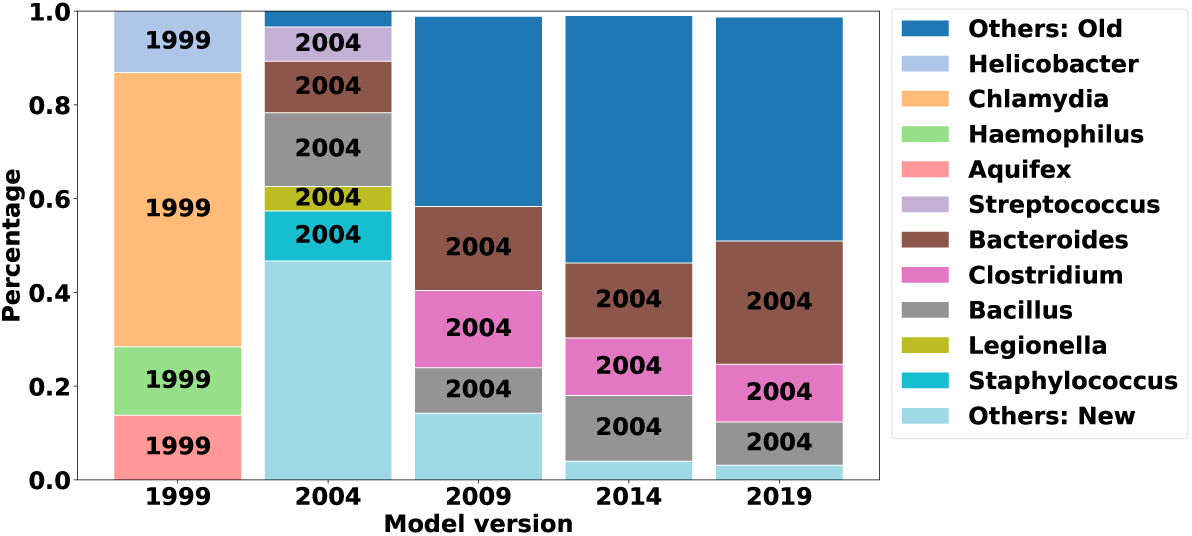
Profiling results change over time on Genus level: NBC Incremental learning classifiers trained on different year are evaluated on a real human fecal sample (SRA ID: SRS105153) on Species level. The figure shows that the predicted composition of the sample changed over time and is influenced by the training data. Taxa with lower than 5% relative abundance is moved to either “Others: New” category or “Others: Old” category depending on weather the taxa is recently added or was added in previous section

**Figure 8.**
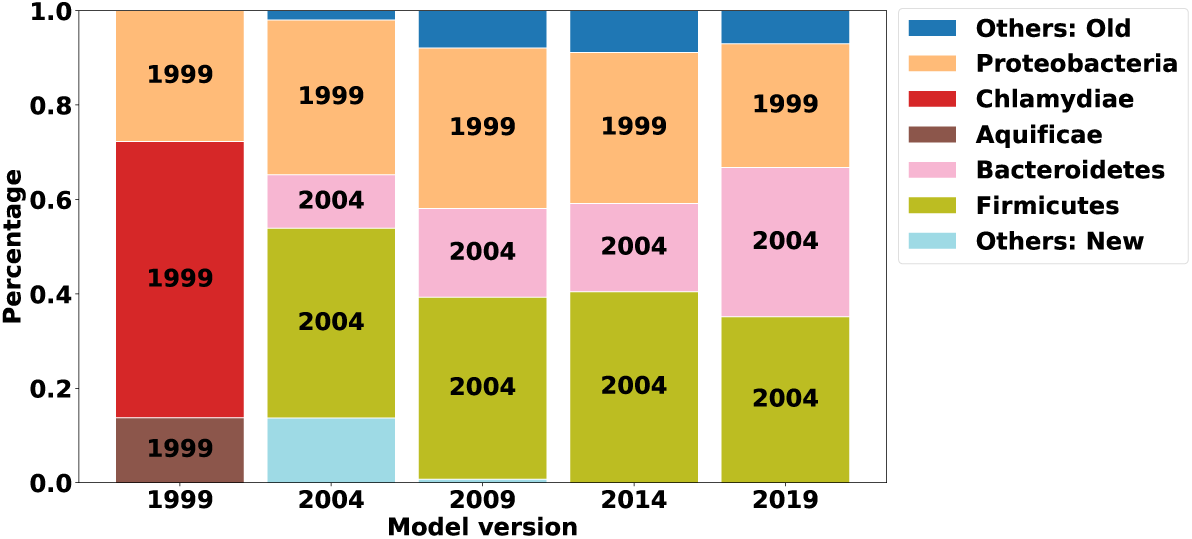
Profiling results change over time on Phylum level: NBC Incremental learning classifiers trained on different year are evaluated on a real human fecal sample (SRA ID: SRS105153) on Phylum level. The figure shows that the predicted composition of the sample changed over time and is influenced by the training data. Taxa with lower than 5% relative abundance is moved to either “Others: New” category or “Others: Old” category depending on weather the taxa is recently added or was added in previous section

As shown in Figures 7 and 8 (and supplementary Figures S7, S8), the profiling results are influenced by the version of training data. For example, in 1999, there are only 4 genomes used to train the database. Therefore, the results in Fig. 8 are drastically skewed and give poor results, with Chlamydiae dominating the gut and no representatives from Bacteroidetes/Firmicutes. As soon as these phylum are introduced by 2004, a great portion of those previously misassigned are assigned to the newly added phyla (and most likely better classifications). We do note that NBC++ is overassigning to Proteobacteria (the amount of Proteobacteria is too high for a healthy gut), due to its over-representation in the training database. However, the percentage of Proteobacteria is steadily decreasing as the amount of training database genomes are getting better, especially with recent additions genomes from Bacteroides/Clostridium/etc. (the proportion being assigned to Bacteroides doesn’t significantly increase until the most recent database version, shown in Fig. 7) that are known to be prevalent in gut samples. NBC’s classifications are biased by training representation, demonstrating the need for up-to-date knowledge to get the most accurate Bacteroidetes/Firmicutes balance.

As we can see on the genus level, in Fig. 7, the amount of classifications being assigned to newly added genera decrease (although more genera are being added now than in the past, see Supplementary file S4_NEW_UNIQUE_GENUS_UPDATES_NUM.pdf in “S4Figure” folder). Similar trends are seen on the species and order levels (see supplementary Figures S7 and S8). After 2009, the top three genera (Fig. 7) in the gut sample remain the same – Bacteroides, Clostridium, and Bacillus (although not always in the same percentage ranking). In fact, it is surprising that the percentage of Bacteroides significantly increases with new knowledge from the past 5 years, demonstrating that there are Bacteroides genomes being added that are very important to classification of this gut. Specifically, on the species level, when Bacillus ovatus is introduced in the 2019 version supplementary Figures S7, many reads get assigned to it, since it is well-known in the gut. However, as described in section “ Mistakes: Current implementation’s tendency to classify to well-represented species classes,” the over-representation of Clostridium botulinum is biasing some of the Clostridium classifications.

### Mistakes: Current implementation’s tendency to classify to well-represented species classes

We note that this NBC implementation has certain defects – which may afflict other classifiers as well. This NBC implementation is biased towards calling class labels that have a large abundance of genomic examples. As a case study, we look at mistakes at the phylum level, arguably the most serious ones, which has a 4% error rate. In the 2019 database, Proteobacteria has 6975 representative genomes while Firmicutes is the next most-represented and has 2925 representatives. Because the algorithm classifies at the species level and then traces up the tree to resolve upper levels, any species that has more occurrences of a *k*-mer as opposed to another species will have an advantage. So, the more genomes that represent a species (or as we sometimes say “examples of that species”), the more likely rare *k*-mers may occur in this particular class (which may accommodate some nucleotide variation).

This biasing of rare *k*-mers can have great impact on the misclassifications that NBC makes, as we can see in the supplementary file `misclassified_reads_table_phylum_level.csv in ‘‘S1Appendix’’ folder`, which is one fold (out of 5) of the phylum level missclassifications. In this fold (1/5 of the entire data), there were 7285 misclassifications on the phylum level out of 186300 reads (96% accuracy). 2663 out of the 7285 (over 36%) misclassifications, tended to classify reads as *Clostridium botulinum* (taxid:1491). This is a problem that becomes even more evident on our test on a real sample (see section “ Evaluation of NBC++ on a healthy gut sample taxonomic profiling results over time”). One outstanding example (in the phylum misclassified spreadsheet) is the Buchnera aphidicola (taxid: 9) species, which had 506 sequences misclassified by NBC, 363 of them classified to *C. botulinum*, 93 classified to *Bacillus cereus/thurigiensis* and another 30 to *C. difficile*, all well-represented species in NCBI. As a reference point, there are 62 genome representatives of *B. aphidicola* while there are 242 genome representatitves of *C. botulinum*, 295 representatives of *C. difficile*, and close to similar amount of *B. cereus/thurigiensis*. What is apparent, is that when some of these sequences are BLASTed (one detailed example in the supplementary material), they share identical stretches of nucleotides up to 18 bases long with essential *C. botulinum* protein sequences that have similarity to essential/fundamental proteins such as ABC transporters, two-component sensor histidine kinases, etc. This shows some important protein sub-domains that are conserved between these organisms. However, due to the single nucleotide mutations that cause these sequences to be unique and prevalence of these subdomains across phyla, these types of sequences are causing NBC to bias its classification towards species that have more representation in the database. This is an important issue to be explored to improve NBC in the future and highlights the importance of keeping up-to-date training data in classifiers and good representation in the database.

## Conclusion

In this paper, we show that new genomes added to the NCBI RefSeq bacterial genome database are exponentially growing. We demonstrate that supervised classification is drastically improving each year with updated NCBI Refseq genome data. By processing simulated reads and a real metagenomic gut sample, we show that new additions to the database change results significantly and have yet to show signs of convergence (i.e. the training database has enough species diversity and example genomes per species to classify samples consistently). Therefore, it is very important to keep the training model updated and that the update process be efficient as possible. We demonstrate that we can achieve efficiency by a proof-of-concept incremental NBC++ taxonomic classifier. By “incrementalizing” taxonomic classifiers, we show that species can be updated/added to the database without the cost of reprocessing the existing database. Our simulation shows that (1) no accuracy is lost compared to the non-incremental NBC implementation and (2) over time, the classification accuracy is a function of the amount of taxa known. Overall, the incremental implementation can gain the valuable information added per year at a fraction of the cost – our example shows that by updating yearly, the 2018 training time would only be 1/4th of the full training time of non-incremental version.

## Materials and methods

### Experimental Dataset

In our experiments, we use the NCBI Reference Sequence (RefSeq) Bacterial genome database and taxonomic tree [2]. Genomes labeled as “complete Genome” in assembly level field and “latest” in version status field are downloaded. Genomes are labeled by their species level label. The genomes are split into 5 folds by random. Testing reads are then simulated from the genomes in 5 folds. 100 reads are simulated from each genome by Grinder [31] with no error. The model will be trained on 4 out of 5 folds and tested on the leftover fold. That is to say, we train on some strains from a species and test on other strains from the same species. In fact, due to this 5-fold cross-validation, some species with smaller amount of genomes will not show up in training process. Therefore, it is marked as unknown if it is not trained on when evaluate the performance of the classifier. All genomes and testing reads are processed by Jellyfish [32] to count the k-mers in each file as naïve Bayes taxonomic classifier takes the abundance of k-mers as features. For our incremental taxonomic classification experiment, we further separate the each fold based on the genomes release year in the assembly summary. Therefore for each year, we have 5 folds and we can train and update the model every year based on which folds the model was originally trained on. For the dynamic of taxonomic profiling experiment, we use all our training genomes as the training data and classify a human fecal sample (SRA ID: SRS105153). We train 5 models in temporal order to show the profiling result change. The initial model is trained by complete Bacteria genomes division 1 (genomes added during 1999), then model 2 is obtained by updating model 1 with training data division 2 (genomes added during 2000 to 2004, model 3 is updated by division 3 (genomes added during 2005 to 2009, followed by model 4 with division 4 (genomes added during 2010 to 2014) and finally model 5 is updated by division 5 (genomes added during 2015 to 2019) from model 4. The composition change on different taxonomic levels are evaluated.

### Naïve Bayes Taxonomic Classifier

Our method (NBC++) is based on a previous non-incremental taxonomic classifier [6]. We then expand the classifier to incrementally update itself efficiently and produce reliable classification results. The core of our classifier is a naïve Bayes (NB) classifier. NB classifier exploits Bayes rule to tackle classification problems which assumes independent features. In essence, NBC’s premise is to maximize the (log)-likelihood of observations given each potential class, 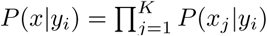, where *x*_*j*_ is the *j*th feature, and *y*_*i*_ is label of *i*th class. The NBC then computes the discriminant function *Q* (a function of the posterior probability) for each class *y*_*i*_ for each observation *x* as *Q*(*y*_*i*_|*x*) = *P*(*y*_*i*_)*P*(*x*|*y*_*i*_), and chooses the class label for which *Q*(*y*_*i*_|*x*) is the largest. In our application, we did not include the prior probability in final likelihood computation because we expect all species equally likely to be observed and don’t know which environment or species prior probabilities to expect. Authors now show that having prior information about species likelihood of occurrence are useful [33]. We compute the probability of observing a k-mer given a species via Laplacian smoothing which defined by the following formula:

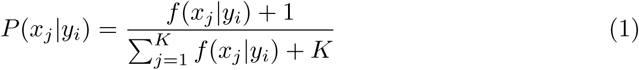

where *f*(*x*_*j*_|*y*_*i*_) is the total number of a k-mer, *x*_*j*_, observed in all training genomes with species level *y*_*i*_. *K* is the total number of unique k-mers for a given size k and *K* = 4^*k*^. The Add-1 smoothing avoids *P*(*x*_*j*_|*y*_*i*_) = 0 when *f* (*x*_*j*_|*y*_*i*_) = 0. Then, for a testing reads, **x**, the posterior probability for a species *y*_*i*_ can be obtained by:

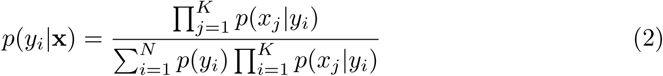

where *x*_*j*_ is one of the k-mer observed in testing reads **x** and *K* is the total number of unique k-mers observed in the testing reads. There are *N* species in the training dataset and *p*(*x*_*j*_ *y*_*i*_) is the probability of observing a k-mer given a species. The prediction is made by taking the argmax of all posterior probabilities. In our implementation, we compute equation 2 in log-space. For example, the numerator of 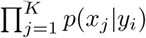 in log space becomes the sum of log-probabilities: 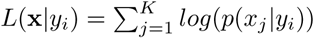.

Traditional NBC implementations require the entire labeled training data to update the model each time when new data are available. Our implementation computes equation 1 only for the added or updated species. The log-space version of equation 2 is still computed for each “read query”.

### Incremental Naïve Bayes Classifier

The incremental implementation of naïve Bayes classifier is quite straightforward. When new sequences are available, we can compute their k-mer of frequency table. Then *P*(*x*_*j*_|*y*_*i*_) is updated based on the label. For example, if a new genome is from one of the existing class, then, the frequency of k-mer *x*_*j*_ is updated and *P*(*x*_*j*_|*y*_*i*_) is updated. If the genome is from a novel class, we create a new class in the model and the *P*(*x*_*j*_|*y*_*i*_) is the frequency of *x*_*j*_ divided by the total number of k-mers. In our implementation, NBC++ performs incremental learning in a number of seamless features that allows it to save on work already performed in the past. By structuring trained models into multiple separate files for each class, new classes can be automatically added when encountered by the training crawler. In this case, updating the model by a new class is as fast as training on that new class alone. The retraining penalty incurred by incorporating new classes into existing models is eliminated in this way. And adding new data to existing classes is also trivial. By storing some partial data alongside pre-computed priors in its save files, NBC++ can seamlessly expand a class in just the time it takes to train on the number of additional reads being appended to the existing database. This eliminates the need to retrain on sequences which are already part of the database, and greatly improves performance when there is a need to frequently update our database with new information. Our algorithm can also be extended to allow taxonomic labels merging in the future. For example, species A is merged to species B in NCBI RefSeq database, we can first update the frequency of *x*_*j,B*_ by 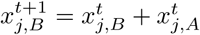 where 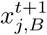 refers to the new frequency of k-mer *j* of species *B*. And then corresponding conditional probabilities can be updated accordingly.

### Scalability of NBC++

In addition to incremental learning, NBC++ introduces a set of improvements meant to optimally leverage the computational resources at its disposal. NBC++ supports multi-threaded operation, making both training and classification faster by parallelizing independent operations such as working on multiple classes or reads simultaneously. By using a command-line parameter (`-t [threads]`), the user can specify the number of threads made available to NBC++. The program will then attempt to run as many parallel instances as threads it has available, and load training/testing data into the available memory for all the threads. The scalability of the previous naïve Bayes classifier is improved. NBC++ can now work within a pre-set memory limit. The program will automatically adjust how many reads it loads into memory, dynamically creating multiple “batches” of various sizes, trying to fit as many as possible in one cycle. This is a sub-optimal but necessary operating mode, as dataset sizes often exceed available RAM. The performance penalty of this process in the case of classification is derived from the repeated iterations through the training database – repeated for each batch. The trade-off is to load and unload the same training classes in/from memory multiple times, because we need to classify multiple batches of reads. We describe the detailed implementation below.

One way in which NBC++ facilitates computational scalability is by automatically separating reads into batches, depending on the amount of memory the user decides to use. By using the `-m [size]` argument, the user can instruct NBC++ to keep all batches under the specified memory cap, thus allowing the user to train and classify large datasets on a variety of systems, getting optimal performance on each run without the need for users to manually adjust batch contents. This is particularly helpful when reads in the database have uneven sizes, which would normally make them difficult to manually add to a batch containing a fixed number of reads.

To demonstrate this behavior, consider a scenario where the program is given 510 sequences (of greatly varying lengths) and the memory is capped to 32GBs. Figure 9 depicts how the program would handle batching in this case – the total size of the dataset of reads is 127GBs, so NBC++ begins loading the first 160 into memory. When it detects the next read would exceed its memory cap, the batch totaling 31.7GBs is released for processing. Once processed, the sequences are unloaded from memory and the process restarts, loading the next batch of sequences until the memory capacity is reached (this time, 50 larger reads take up 31.9GBs). Note that class savefiles will also be loaded for each thread in addition to the loaded reads, therefore the memory cap is only used to dynamically adjust batch sizes.

**Figure 9.**
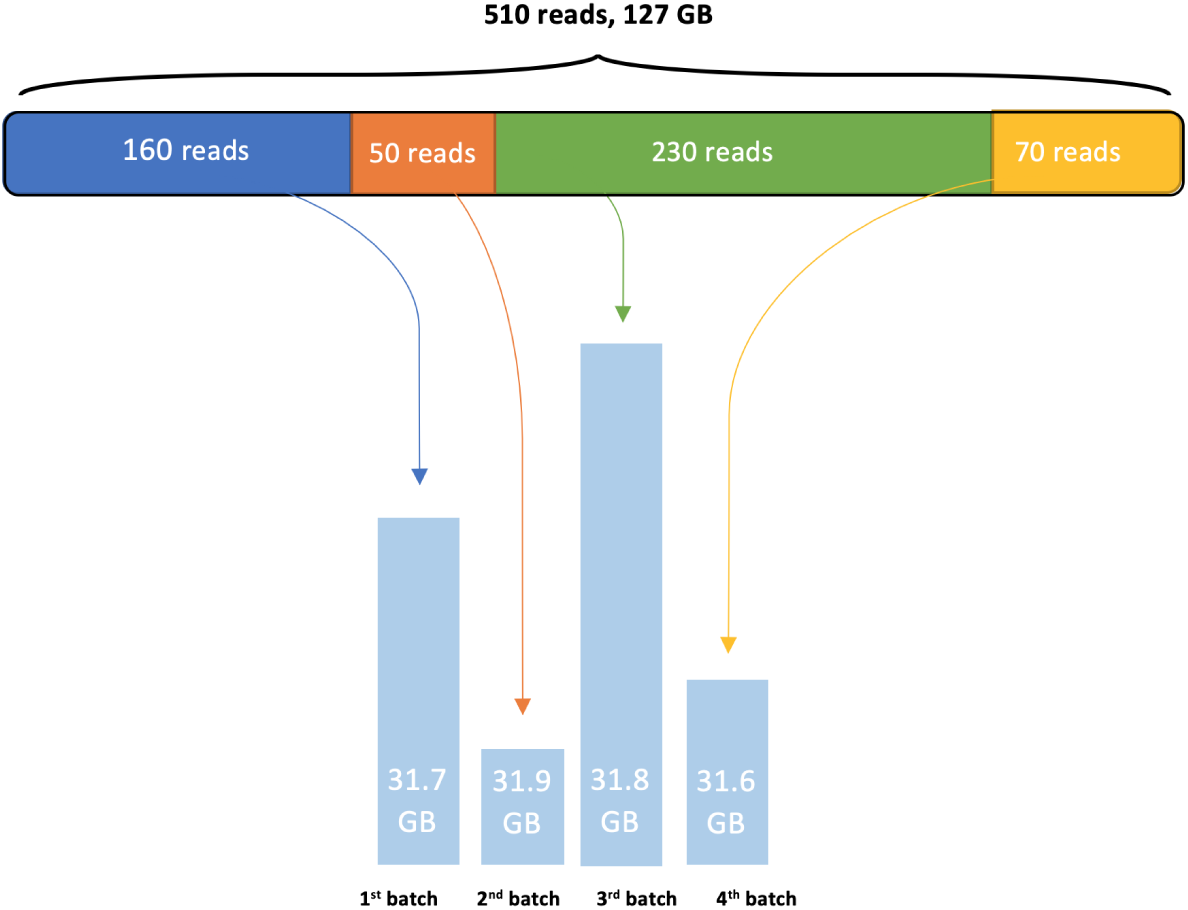
Batch loading diagram. NBC++ handle large dataset by load a batch/subset of data at a time to process. Due to the incremental learning capability of the program, it can easily handle large dataset in both training and testing phases

An additional argument, `-n [nbatch]` is provided as a manual override for the number of reads to include in a batch. This argument is not compatible with the memory cap option and should only be used if the user prefers to process a fixed number of reads at a time, rather than trying to use all available memory optimally.

We also allow for multi-threaded computation through the `-t [threads]` argument, which prompts NBC++ to create worker threads that will concurrently handle one class each. This option is independent of the memory cap, which means that users should allow for some additional memory to be used by each thread in loading their training class savefiles: `total_memory = read_memory_cap + n_threads × savefile_average_size`.

With these two options enabled, NBC++ will first create a batch of reads that fit within the provided memory cap. The program will then create the specified number of threads, loading one class savefile for each and processing in parallel all reads within the batch. When we’ve finished processing all the reads in the batch, the class savefile is unloaded and the next class is read from disk. The thread will then begin to process reads from the beginning. The batch is discarded when this operation has ran for all existing classes, in which case the program will begin creating a new batch and repeating the process until the entire dataset is processed.

The multi-threaded computation along with memory cap is very useful in computing cluster environment. We use a case study here to illustrate the usage. Suppose we allocated 16 cores and a total of 64 GB memory resources to run NBC++. Then, the value we pass to `-t [threads]` would be 15 since we need 1 core for master process and the rest of 15 can be used for multi-threading. The memory cap should be considered as the memory available solely for testing reads. Given we have 16 cores and 64 GB memory, we have 4 GB per core. If the maximum size of training class savefile (class object of a species derived from training data) is 2 GB. Then 2 GB per core are available for testing. To be safe, we don’t include the memory for master core for testing reads. Then the total amount of memory for testing can be set to 2 × 15 = 30 GB. Therefore, we pass 30 GB to `-m [size]` parameter. In this way, we manage to balance the loading of both save files (training) and testing data in memory.

## Supporting Information

All supporting information can be found in the supplementary files. Here we provide the details of the supplementary files.

**S1 Appendix. Case example of B. aphidicola and C. botulinum**

Take the following read as an example:

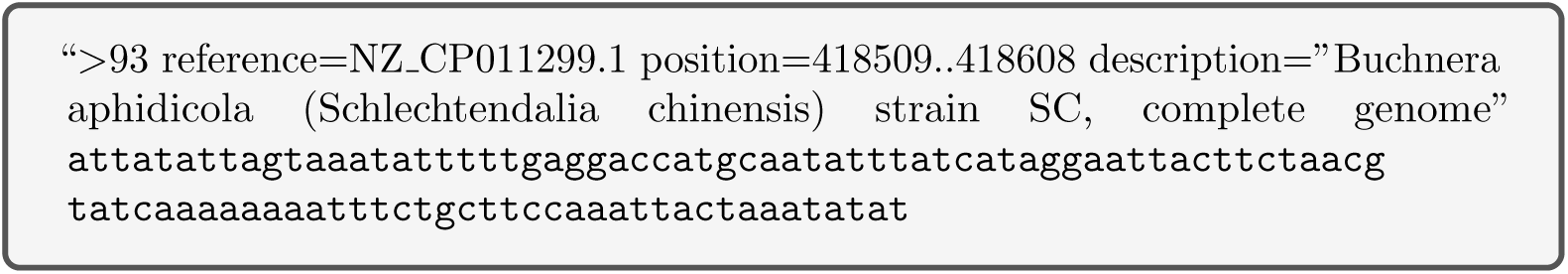

This sequence shares 11 15-mers with the *B. aphidiciola* (taxid:9) that were trained on for the first fold (see supplementary file `kmr_9.csv` in “S1Appendix” folder), while it shares 18 15-mers with *C. botulinum* (taxid:1491) (see supplementary file `kmr_1491.csv` in “S1Appendix” folder). While many of the 18 k-mers in *C. botulinum* had lower probability accounting for the fact that *C. botulinum* had 13× the amount of nucleotides in its class than *B. aphidiciola*, the fact that 7 more unique *k*-mers were found, was substantial enough to tip the classification towards *C. botulinum*. As a comparison, the *B. aphidicola Schlechtendalia chinensis* sequence was blasted against other *B. aphidicola* strains and also against *C. botulinum*. The queries match to the *B. aphidicola* sequence with similarities of (∼58/59% for long stretches and 80-90% identities for short stretches), for other strains of *B. aphidicola* excluding strain Schlechtendalia chinensis in supplementary file `blast_118110_against_other_9_no_best_match.pdf` in “S1Appendix” folder. Similar similarities are found for the blasts of the *B. aphidicola* query against *C. botulinum* seen in supplementary file `blast_118110_against_1491.pdf` in “S1Appendix” folder. (If the PDFs only yield an incomplete picture for the reader, we also have search strategies, ending in .ASN in “S1Appendix” folder available to explore this blast). The BLAST results show that even BLAST has slight difficulty distinguishing these sequences.

**S2 Appendix. Experimental data source**

The bacterial genome data is downloaded from the NCBI website according to the GCF assembly summary file as it is in March 2nd 2019 download from ftp://ftp.ncbi.nlm.nih.gov/genomes/refseq/bacteria/assembly_summary.txt. The taxonomic tree is downloaded from ftp://ftp.ncbi.nlm.nih.gov/pub/taxonomy/taxdump.tar.gz as it is in March 2nd 2019.

**S3 Table. New Refseq Bacterial genomes published every year**: the table shows the name and taxid of completed genomes published every year. See “S3TABLE” folder.

**S4 Fig. Number of updates in the NCBI bacteria genome database on six taxonomic level, namely, Species, Genus, Family, Order, Class, Phylum** We have a figure for “Accumulative number of updates per year” per taxonomic level and a figure for “compared with last year, the number of new updates per year” per taxonomic level. See “S4Figure” folder.

**S5 Fig. Species- and Order-Level Accuracy for NCBI example**: The figures show the species and order level accuracy from 1999 to 2019. See “S5Figure” folder.

**S6 Fig. Histogram of number of genomes in Species**: The figures show yearly histograms of the number of genomes per species from 1999 to 2019. See “S6Figure” folder.

**S7 Fig. Profiling results change over time on Species level:** NBC incremental learning classifiers trained on different year are evaluated on a real human fecal sample (SRA ID: SRS105153) on Species level. The figure shows that the predicted composition of the sample changed over time and is influenced by the training data. Taxa with lower than 5% relative abundance is moved to either “Others: New” category or “Others: Old” category depending on weather the taxa is recently added or was added in previous section

**S8 Fig. Profiling results change over time on Order level:** NBC incremental learning classifiers trained on different year are evaluated on a real human fecal sample (SRA ID: SRS105153) on Order level. The figure shows that the predicted composition of the sample changed over time and is influenced by the training data. Taxa with lower than 5% relative abundance is moved to either “Others: New” category or “Others: Old” category depending on weather the taxa is recently added or was added in previous section

## Authors’ contributions

ZZ: Formal analysis, Methodology, Validation, Visualization, Supervision, Writing – original draft & review & editing; AC: Software Development, Formal analysis, Methodology, Validation, Writing – original draft; GR: Conceptualization, Funding acquisition, Resources, Supervision, Writing – original draft & review & editing.

## Acknowledgments

This project is sponsored by CVDI and Becton, Dickinson and Company by NSF I/UCRC grant #1650431. Work reported here was run on hardware supported by Drexel’s University Research Computing Facility (URCF). This work also used the Extreme Science and Engineering Discovery Environment (XSEDE) [34], which is supported by National Science Foundation grant number ACI-1548562.

## Footnotes

1 ftp://ftp.ncbi.nlm.nih.gov/genomes/refseq/bacteria/assembly_summary.txt

2 https://blast.ncbi.nlm.nih.gov/Blast.cgi?PAGE_TYPE=BlastDocs&DOC_TYPE=Download

3 https://aws.amazon.com/ec2/pricing/on-demand/

## Notes

#### Summary of Updates

Section on Abstract, Introduction, Results and Methods are updated to clarify and make our points more clear.

https://github.com/EESI/inbc-supporting-information

