## Supplementary figures and images for "Keeping up with the genomes: efficient learning of our increasing knowledge of the tree of life" (from another version of this preprint: version 1, 2019-09-08)

### 1999.pdf

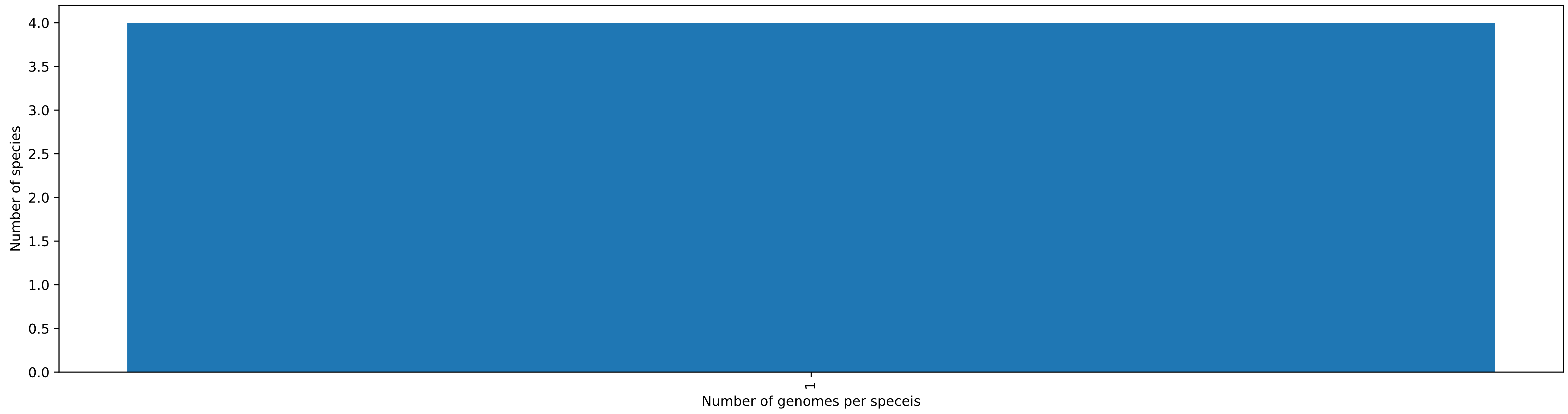

### 1999.pdf

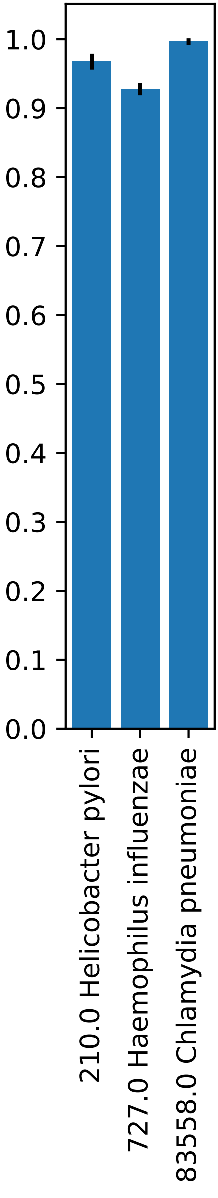

### 2000.pdf

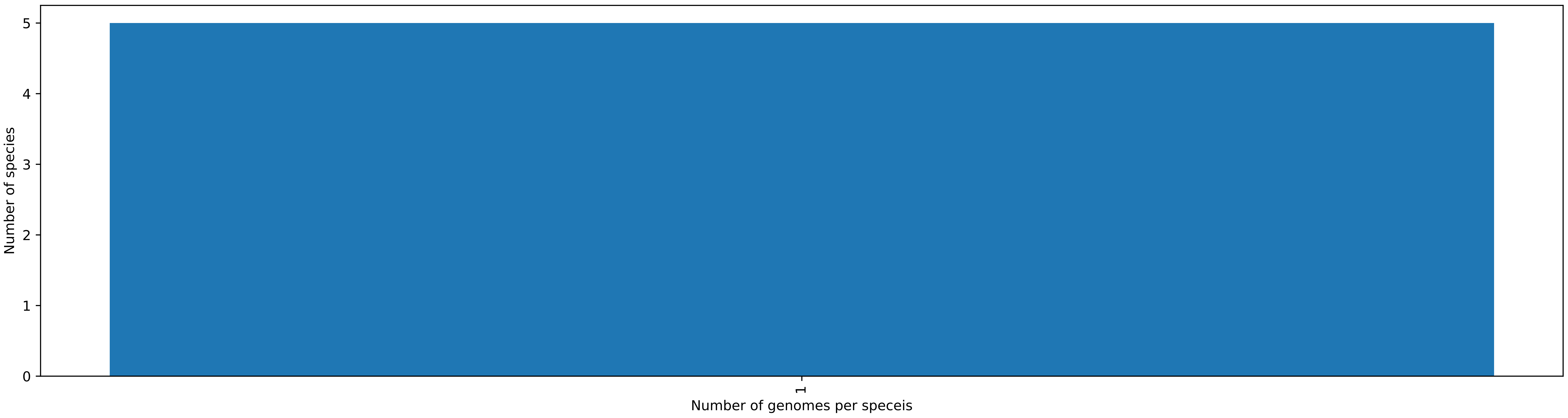

### 2000.pdf

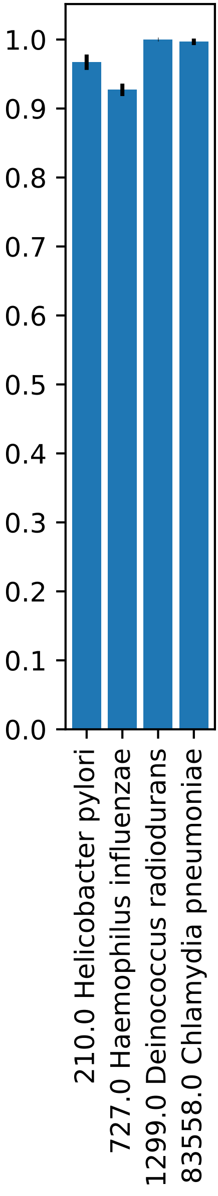

### 2001.pdf

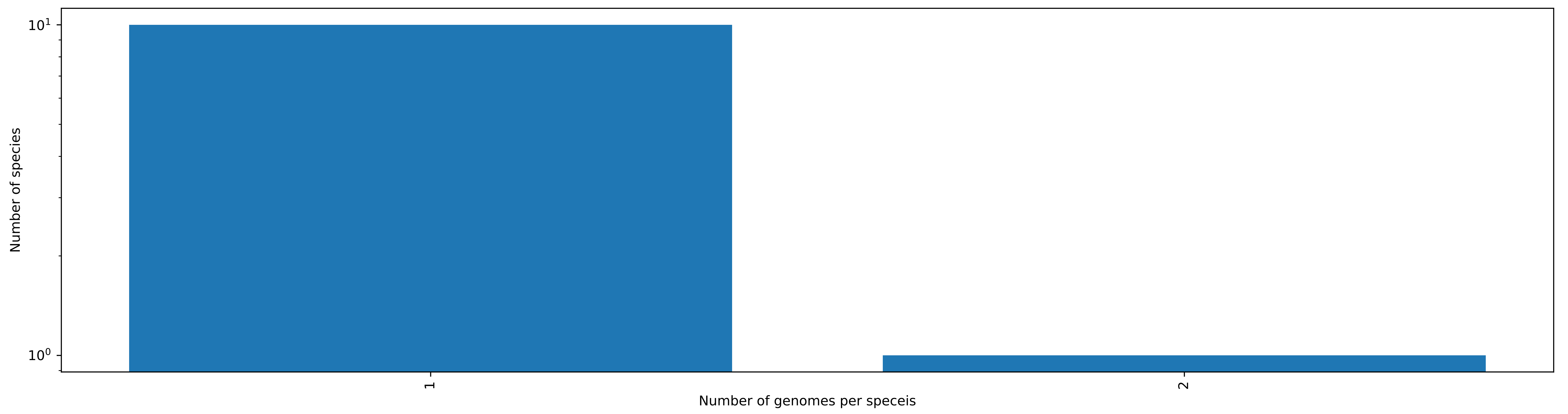

### 2001.pdf

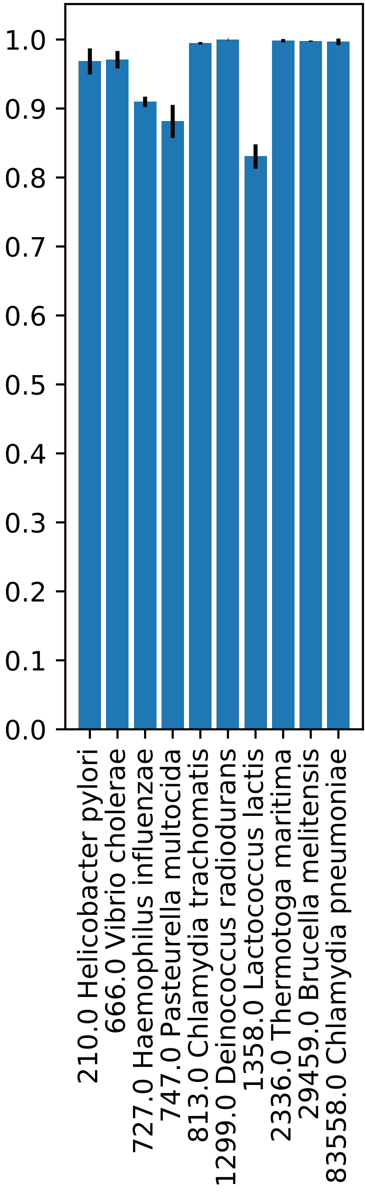

### 2002.pdf

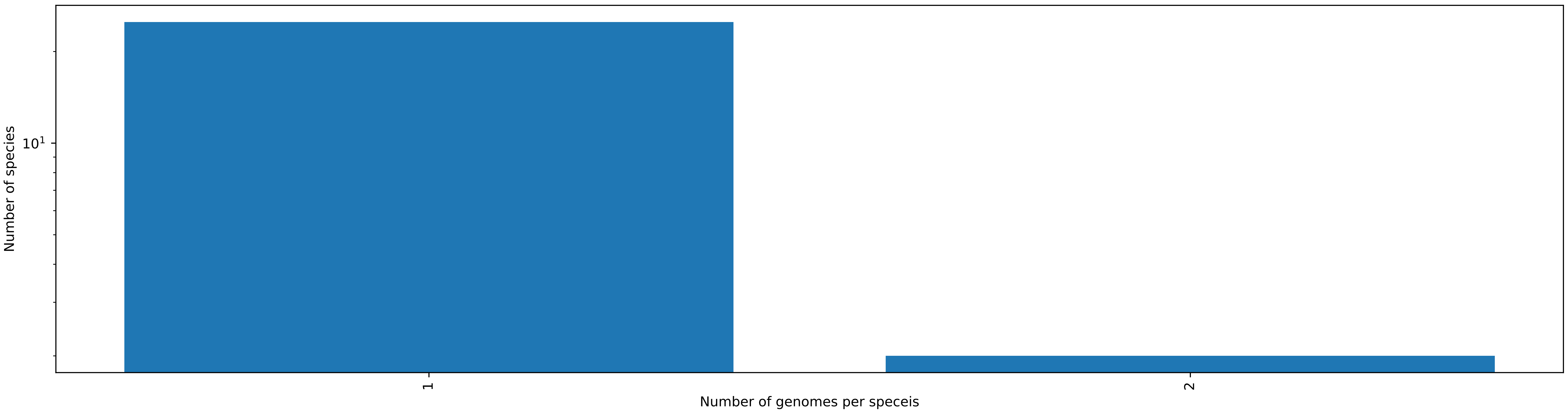

### 2002.pdf

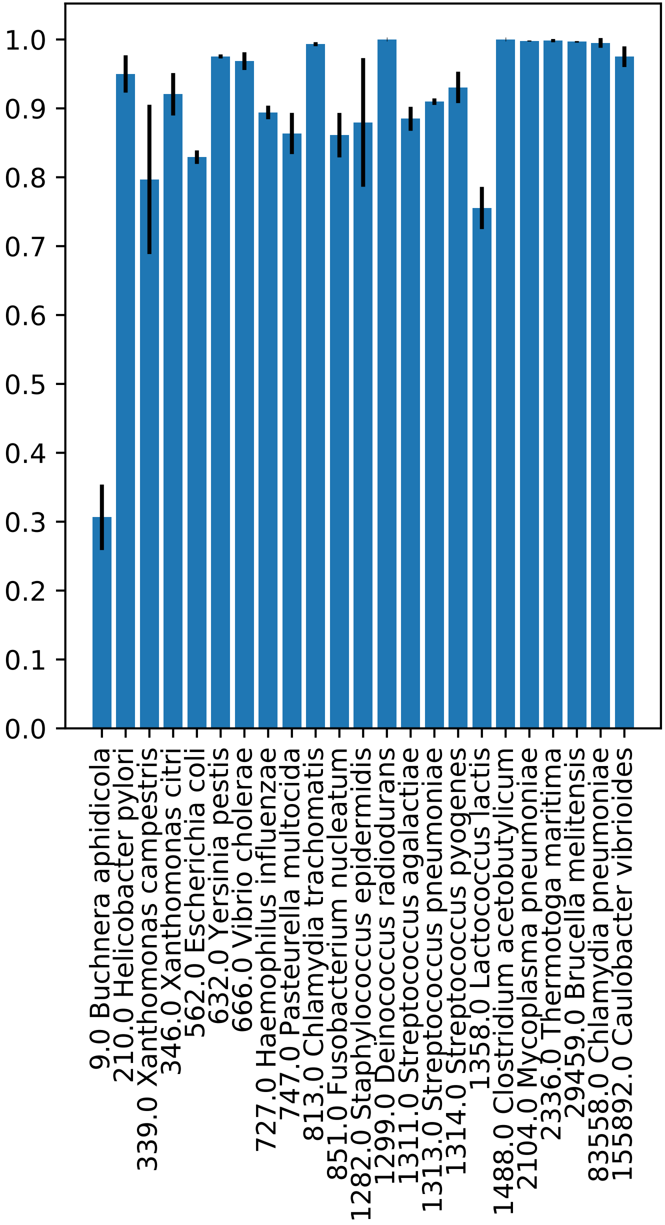

### 2003.pdf

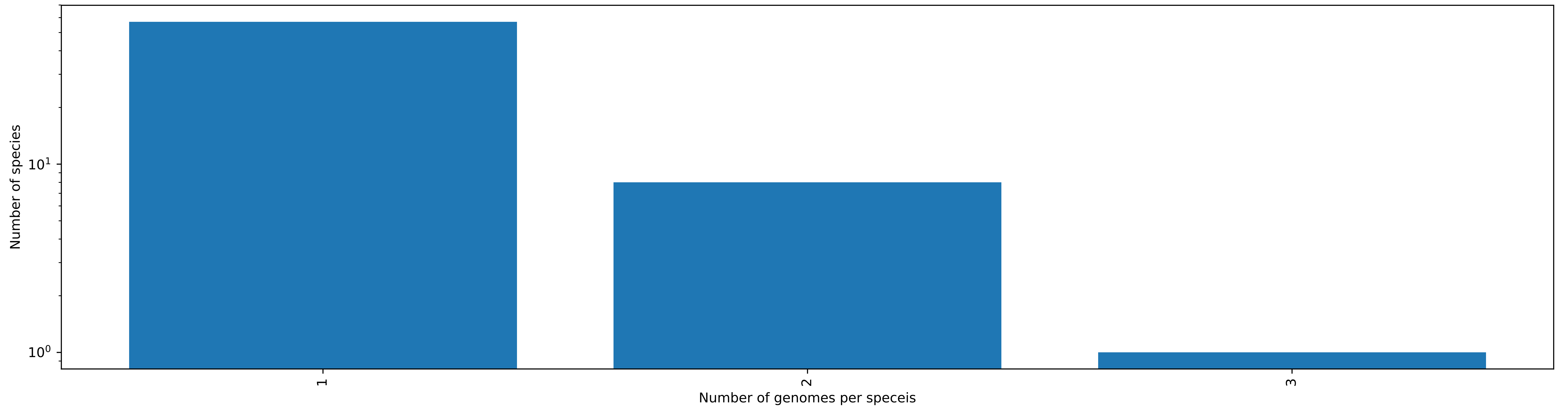

### 2003.pdf

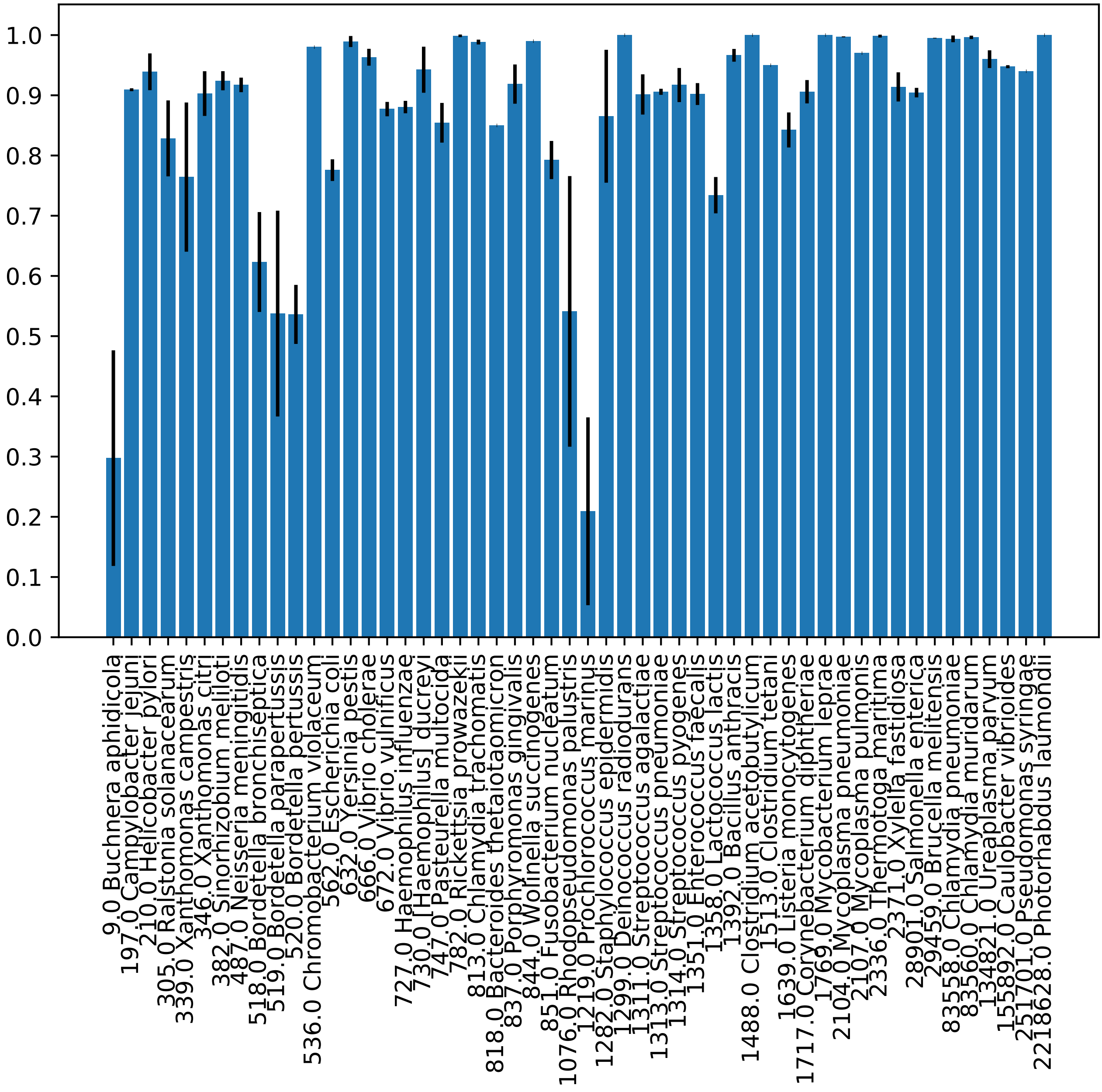

### 2004.pdf

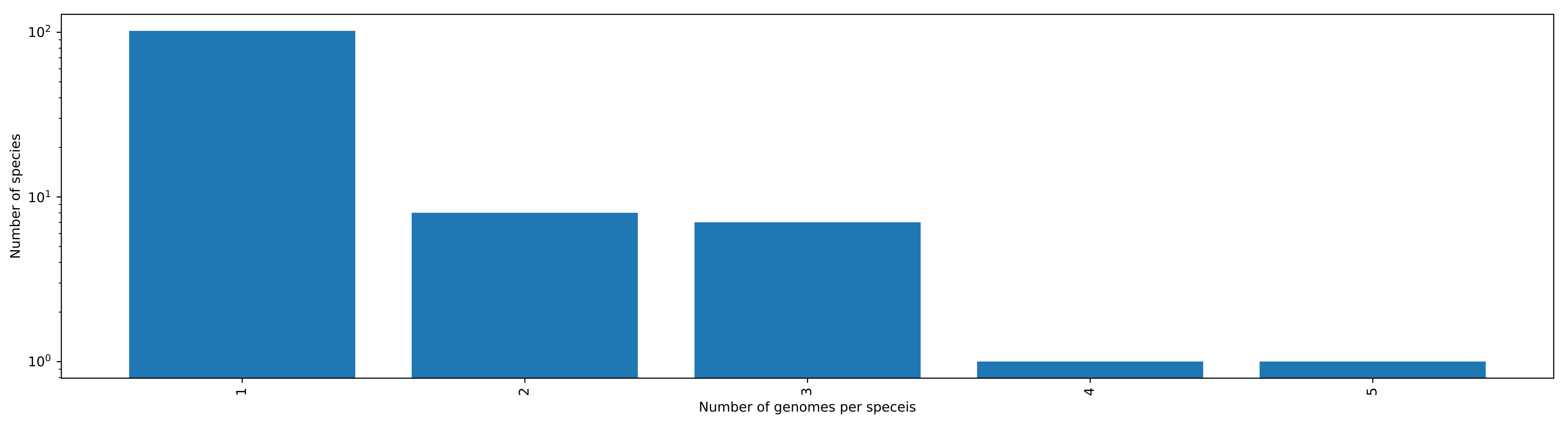

### 2004.pdf

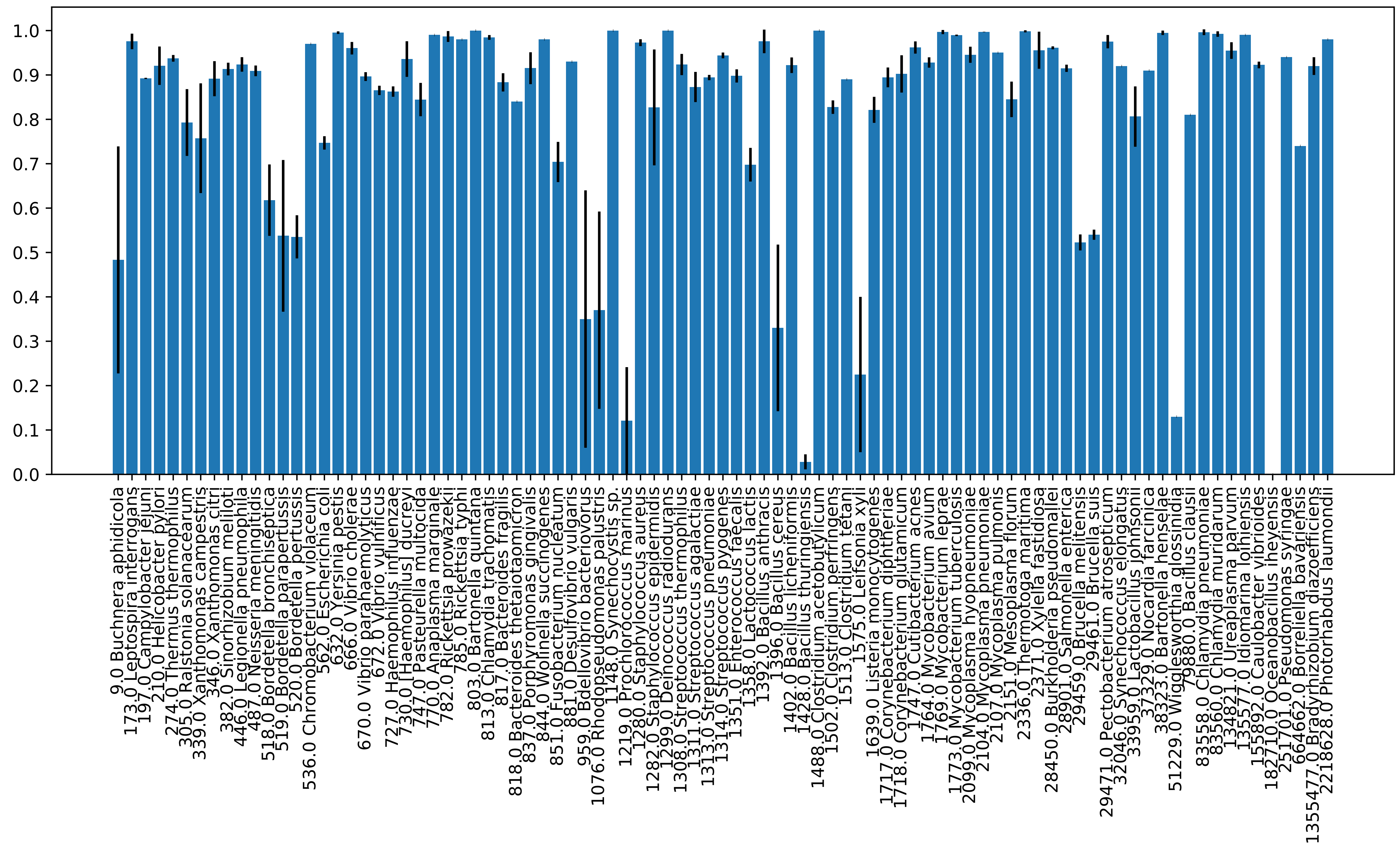

### 2005.pdf

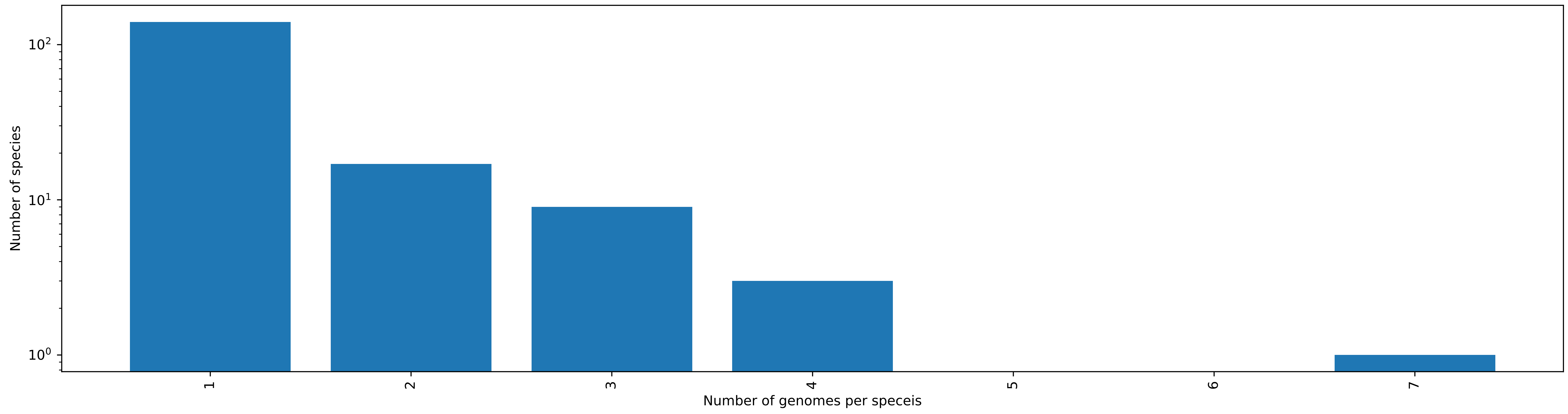

### 2005.pdf

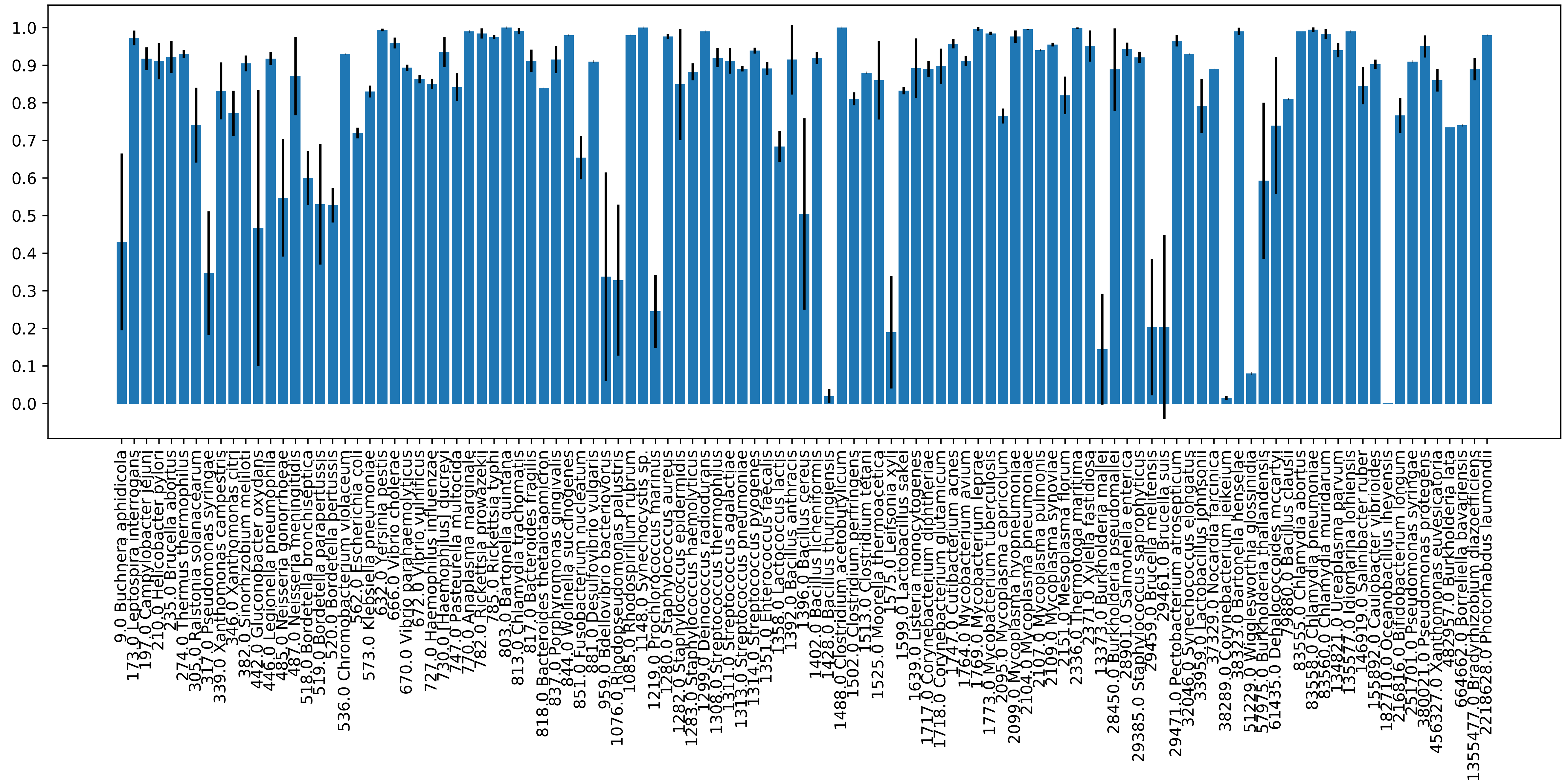

### 2006.pdf

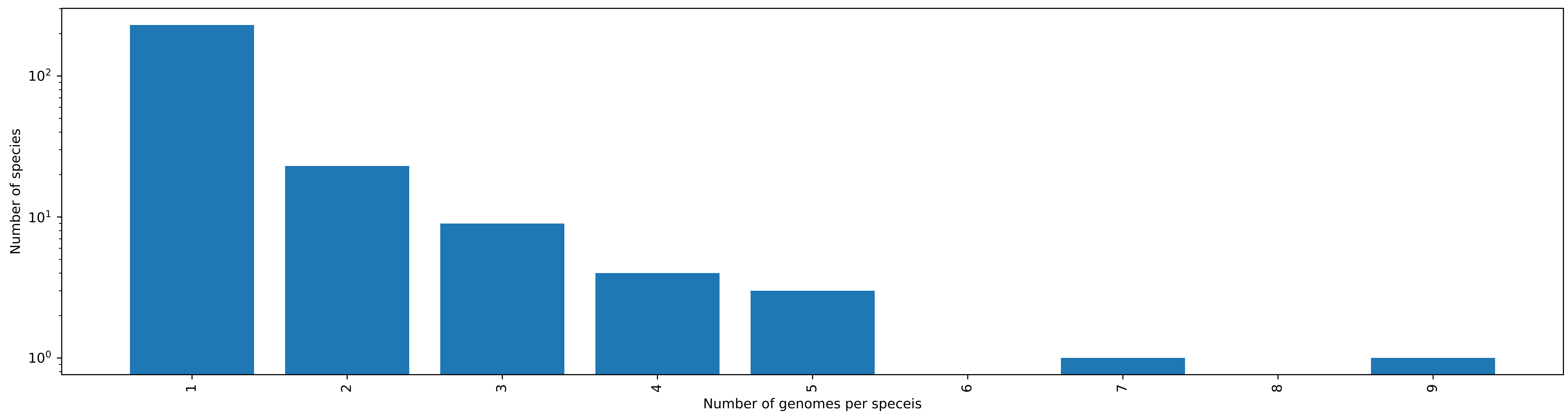

### 2006.pdf

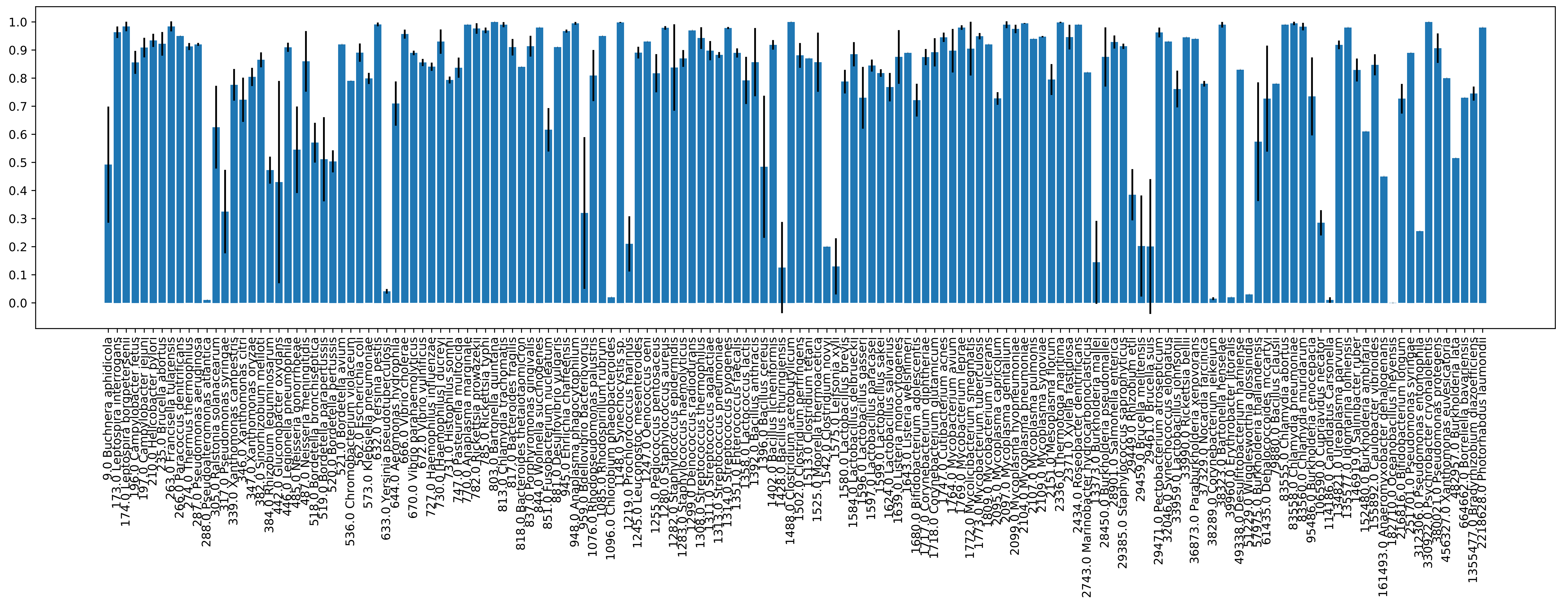

### 2007.pdf

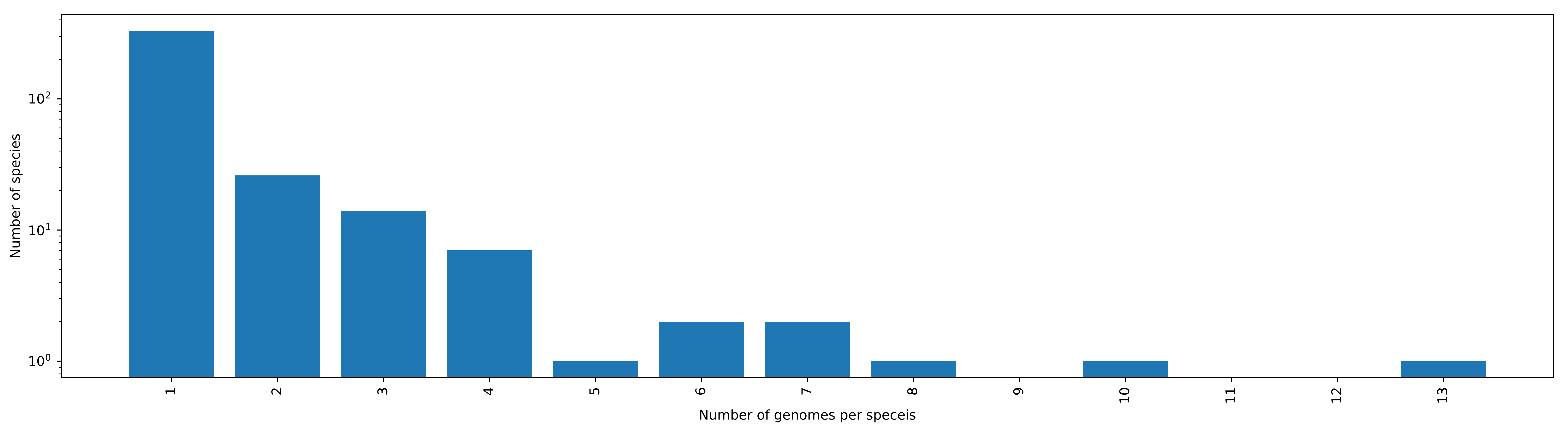

### 2007.pdf

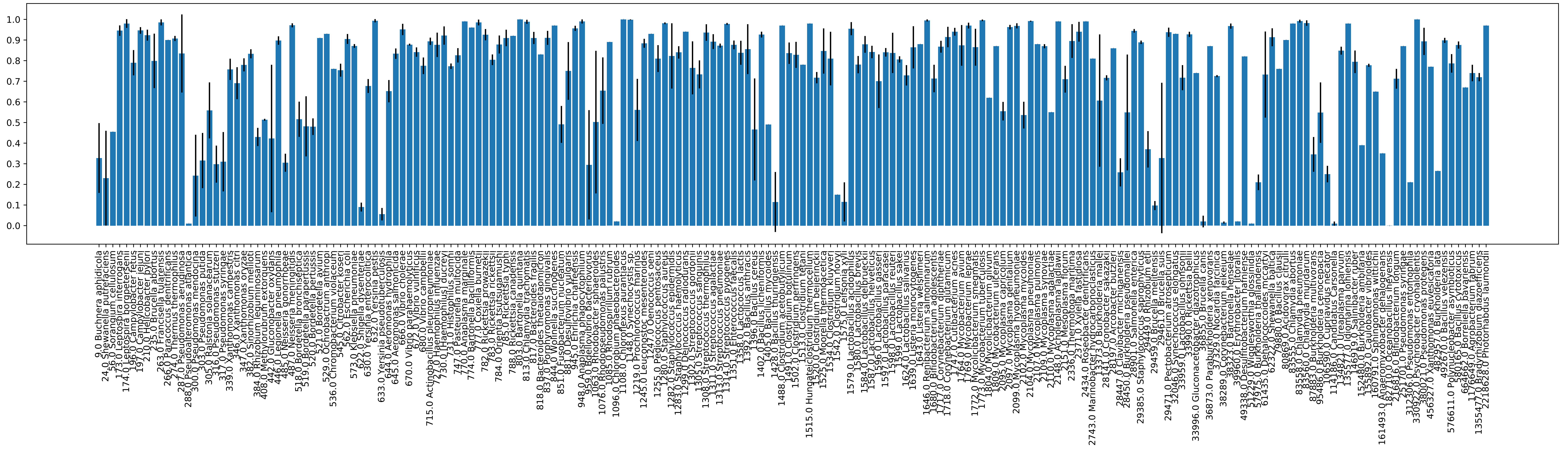

### 2008.pdf

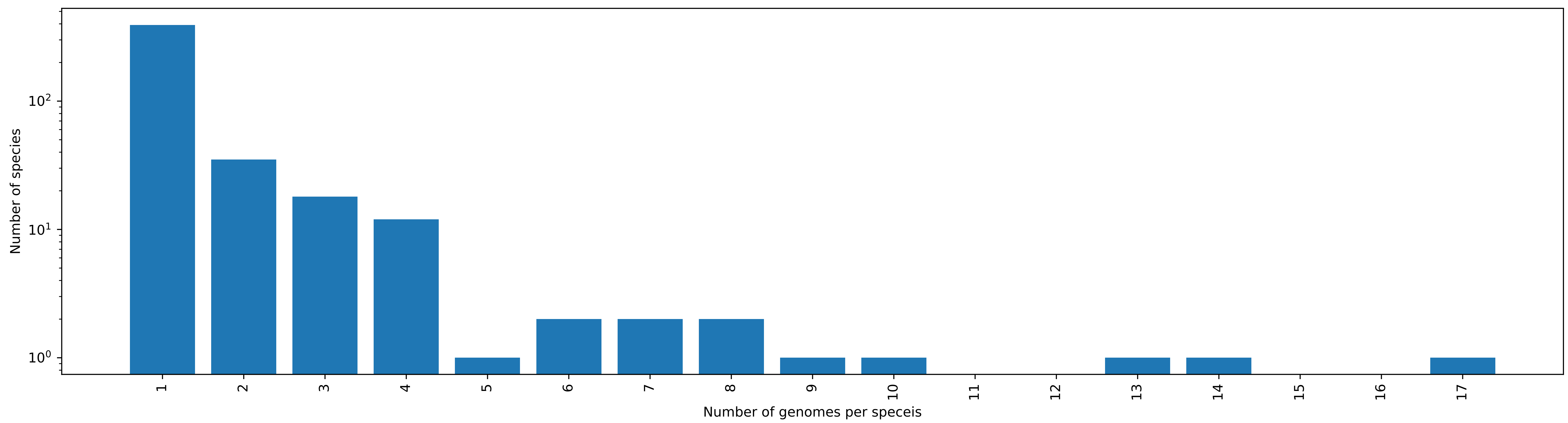

### 2008.pdf

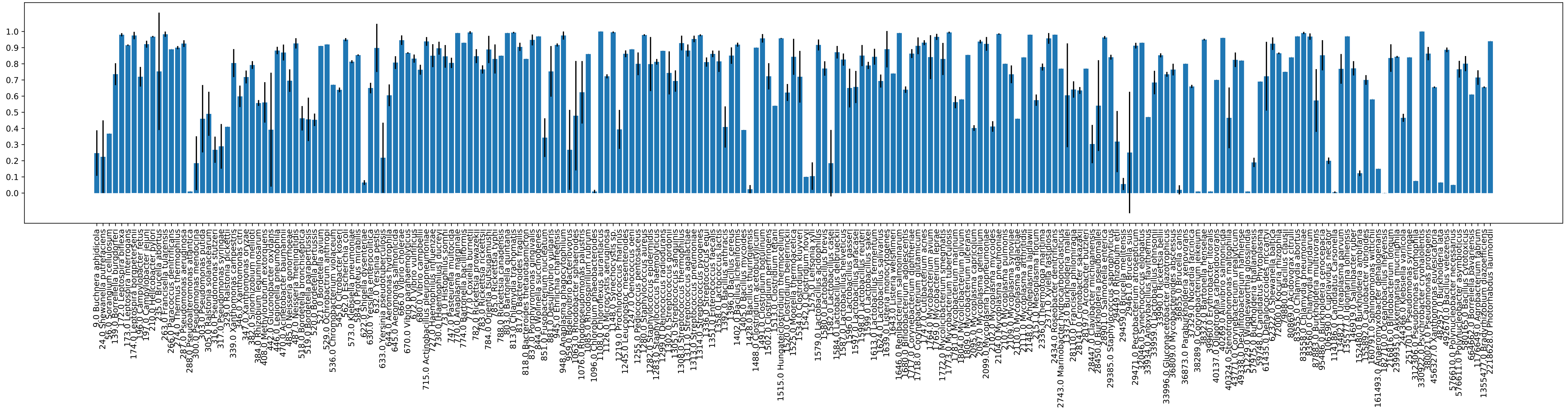

### 2009.pdf

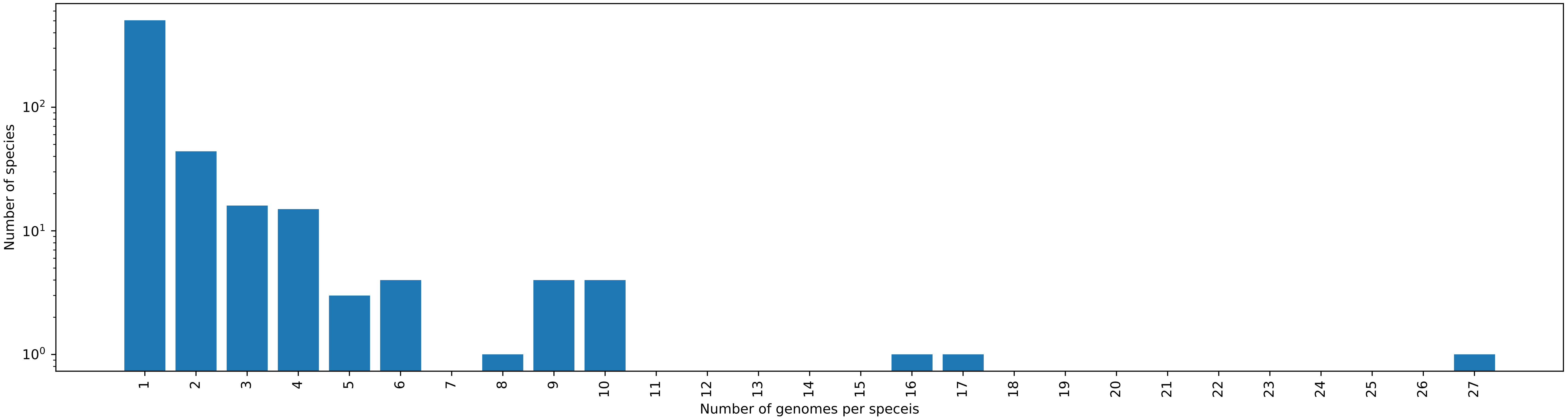

### 2009.pdf

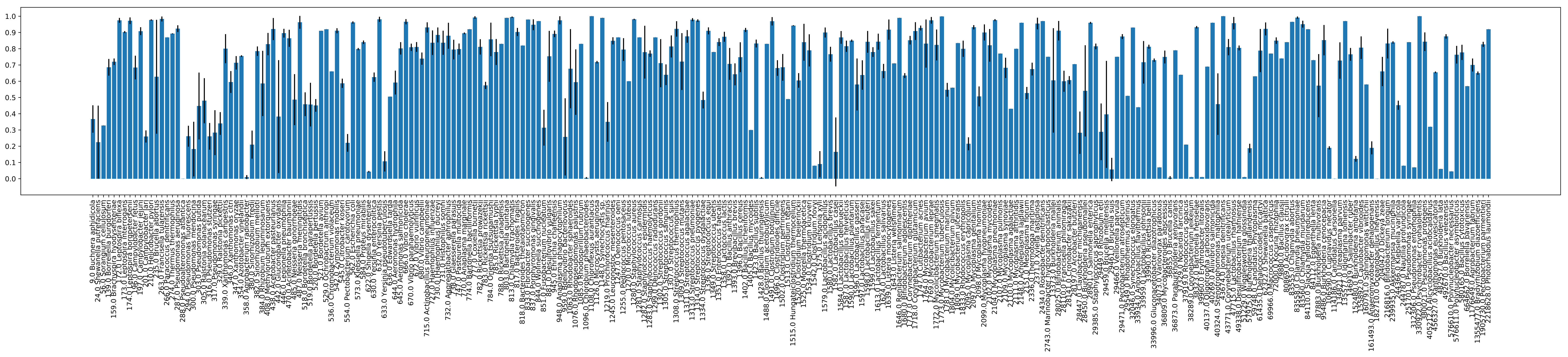

### 2010.pdf

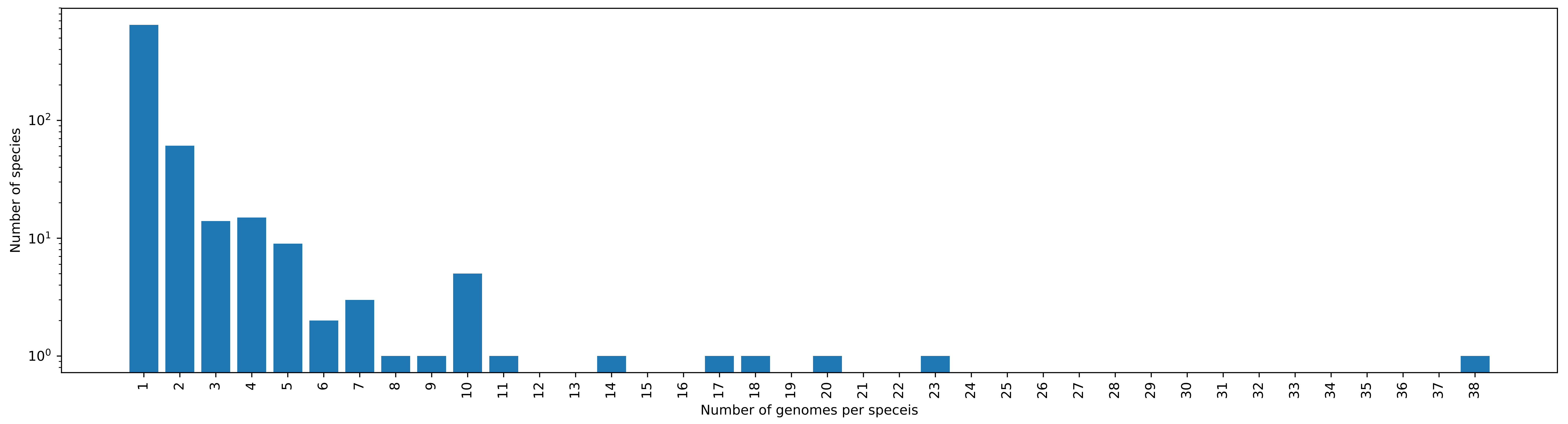

### 2010.pdf

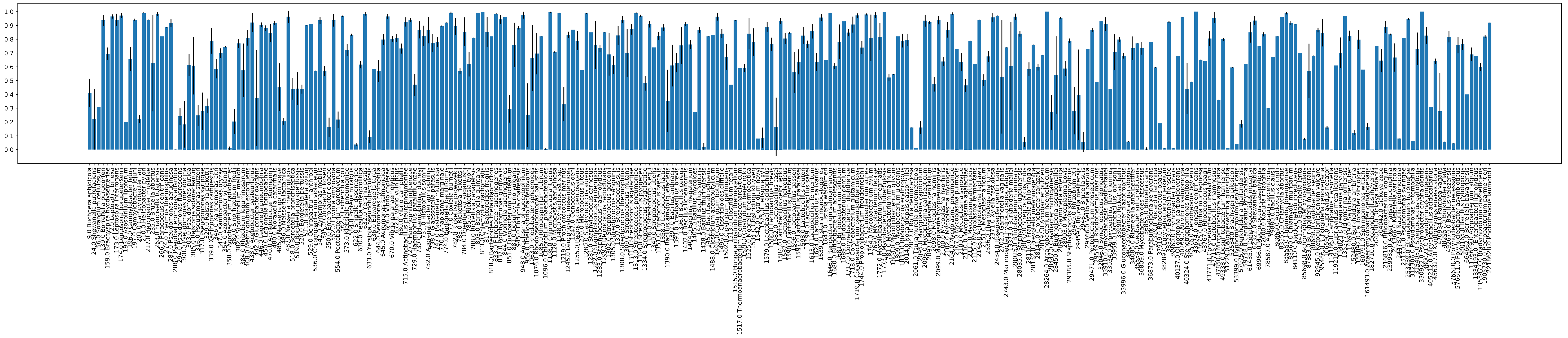

### 2011.pdf

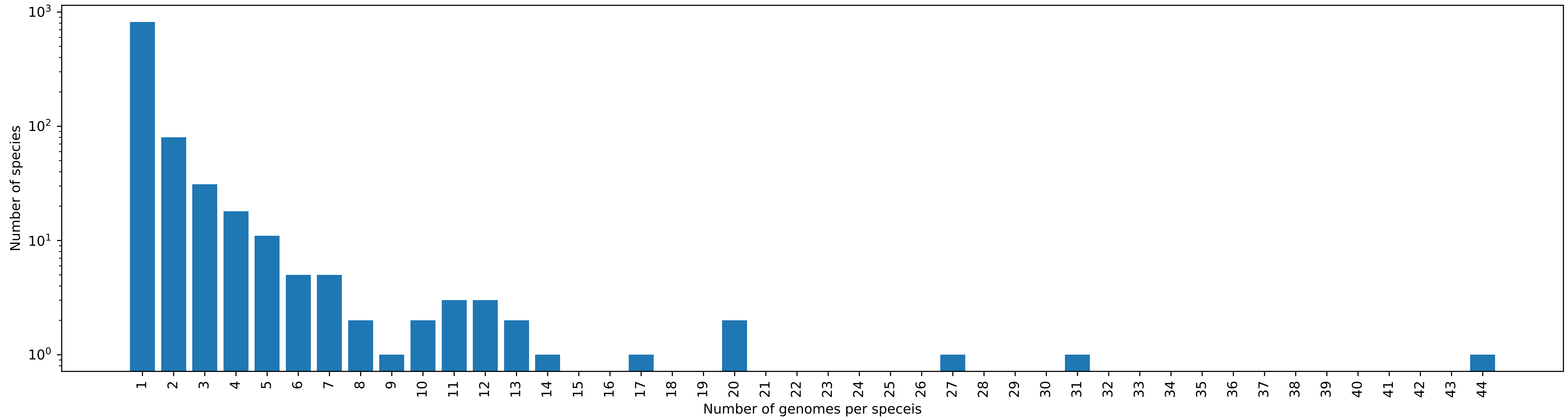

### 2011.pdf

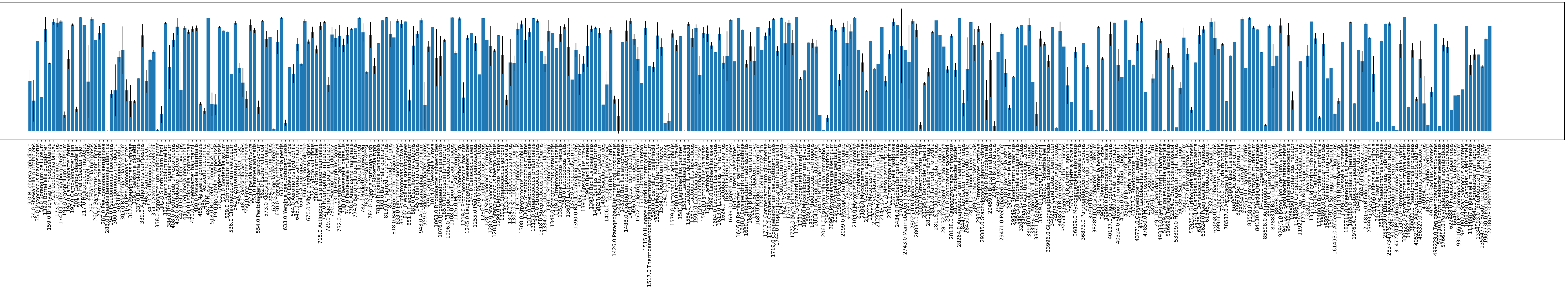

### 2012.pdf

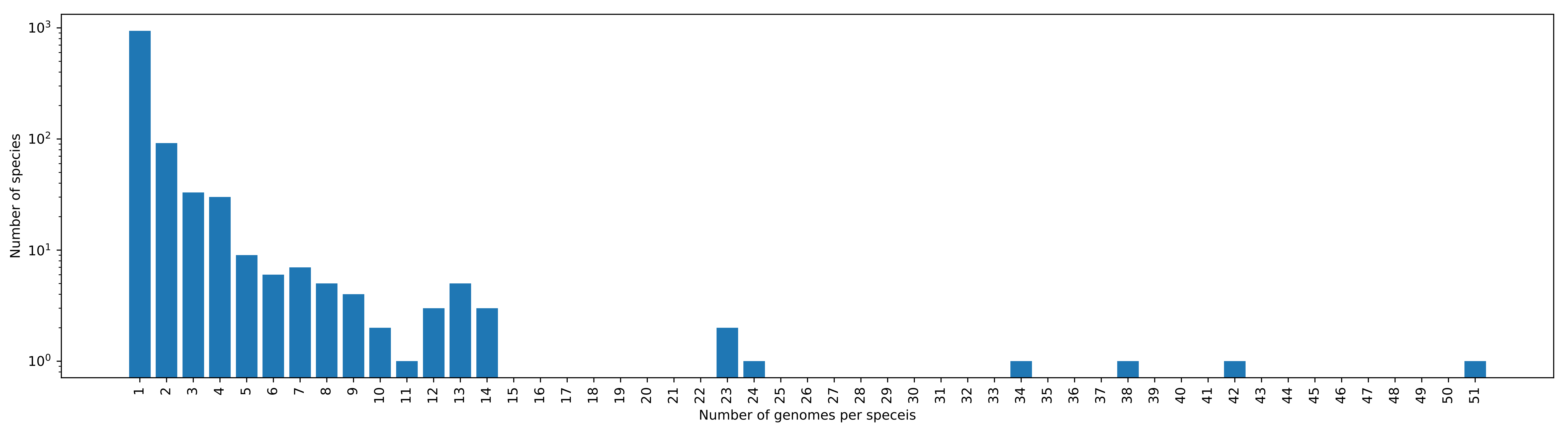

### 2012.pdf

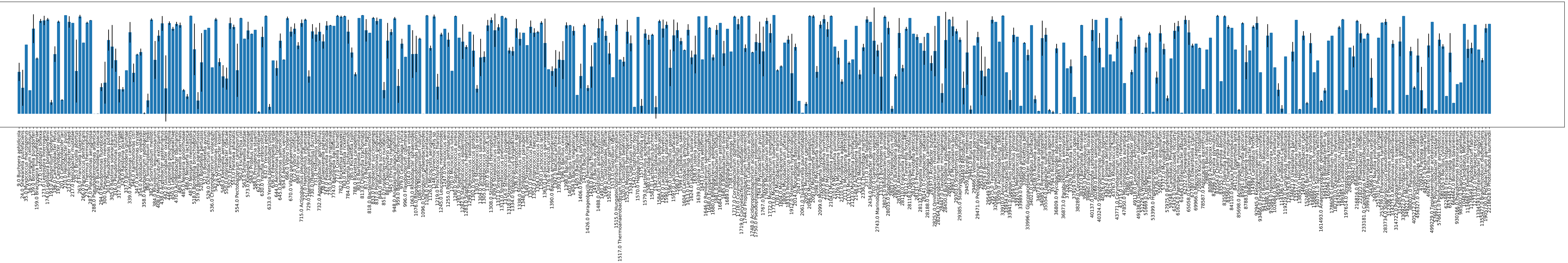

### 2013.pdf

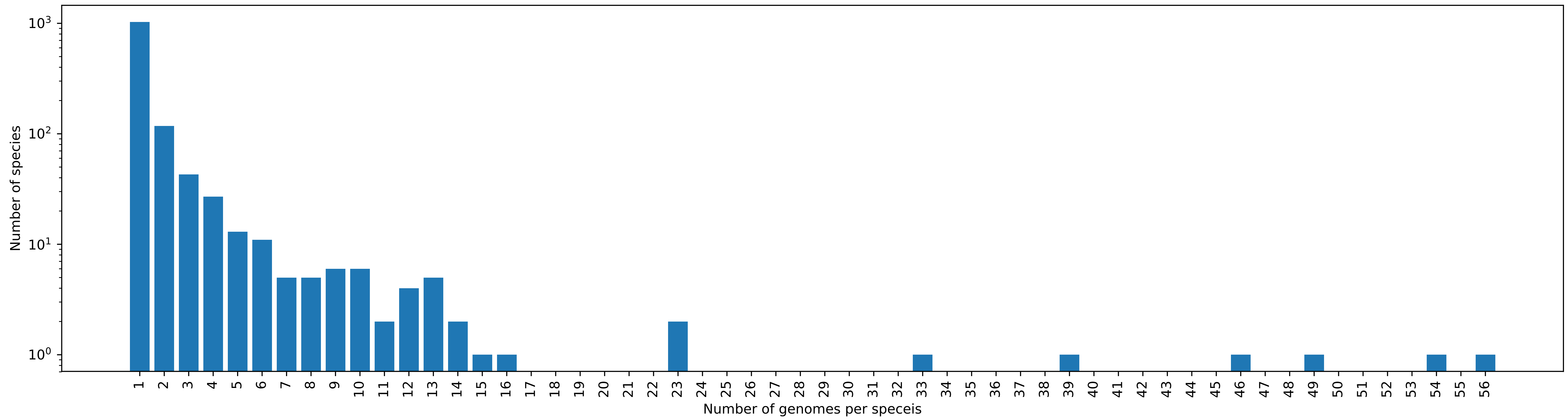

### 2013.pdf

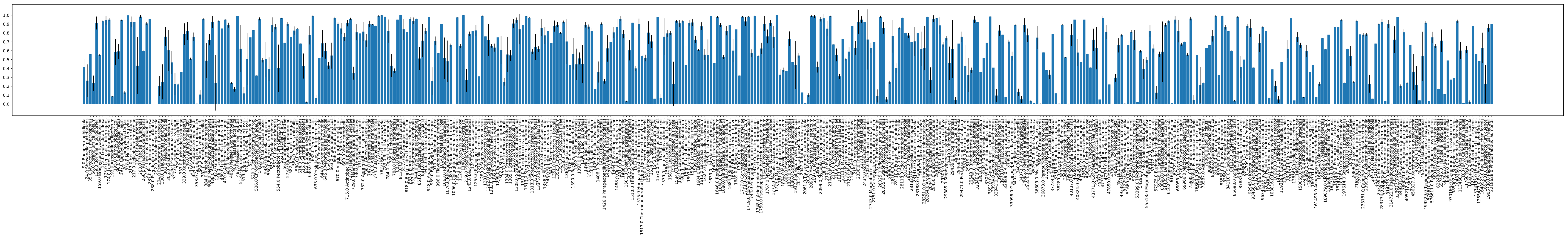

### 2014.pdf

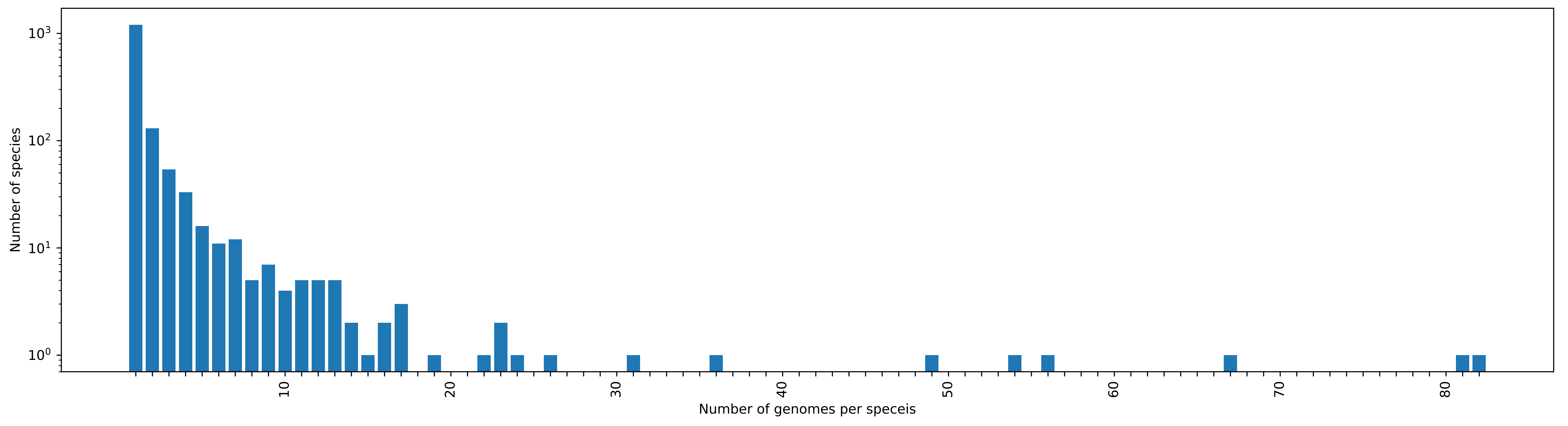

### 2014.pdf

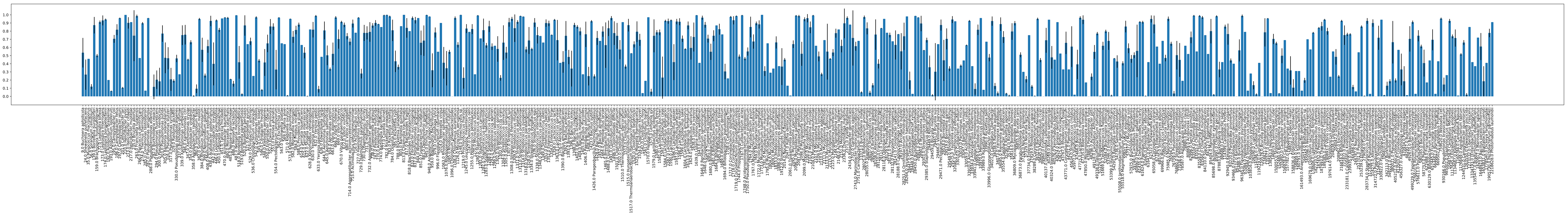

### 2015.pdf

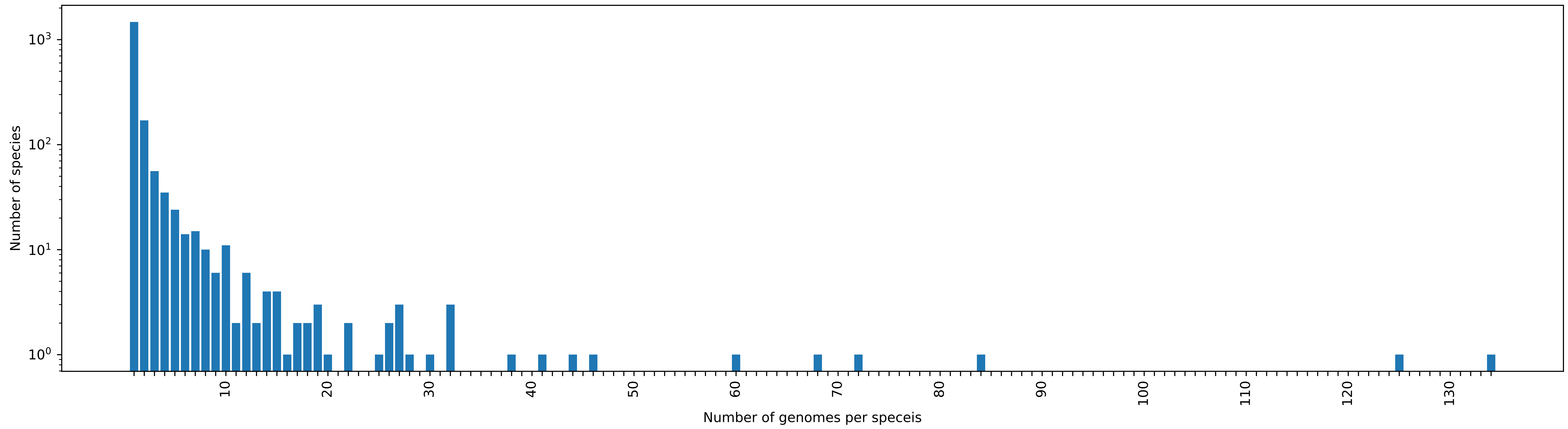

### 2015.pdf

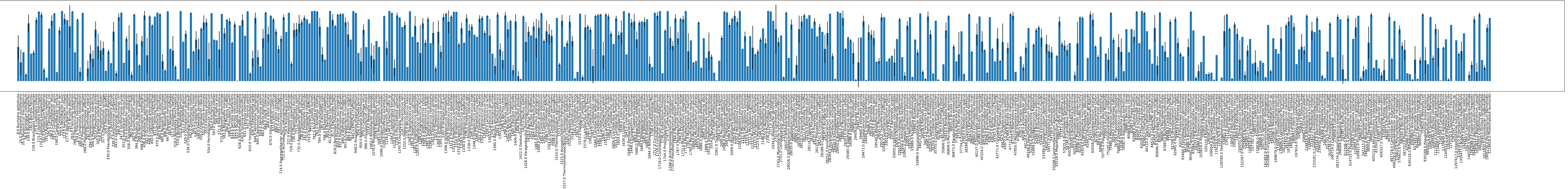

### 2016.pdf

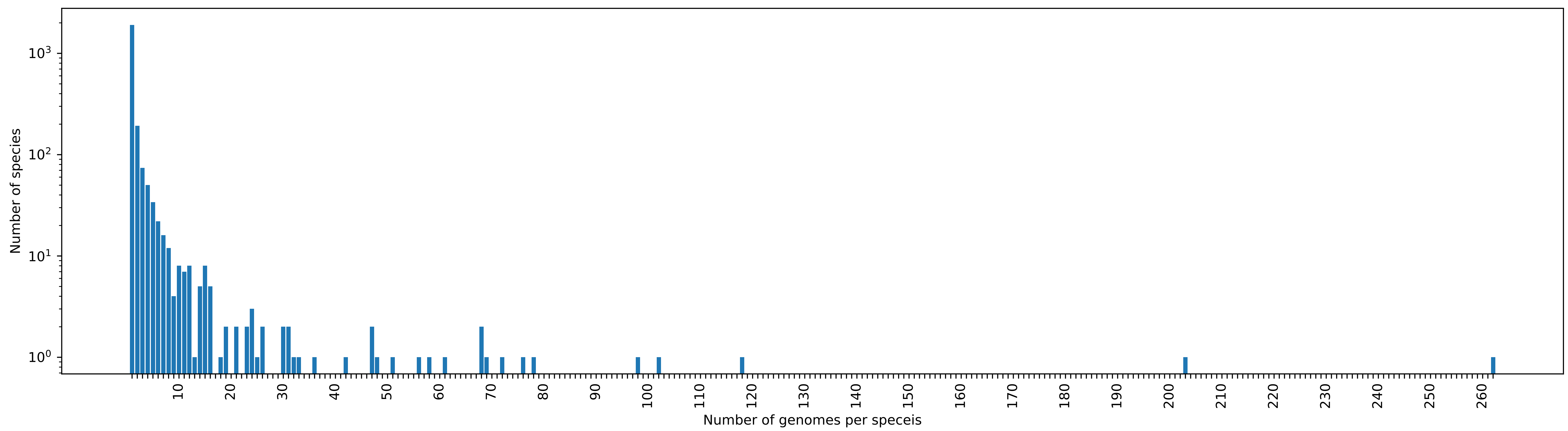

### 2016.pdf

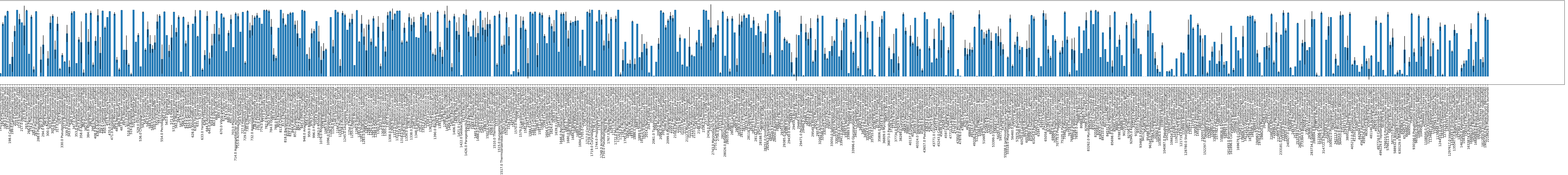

### 2017.pdf

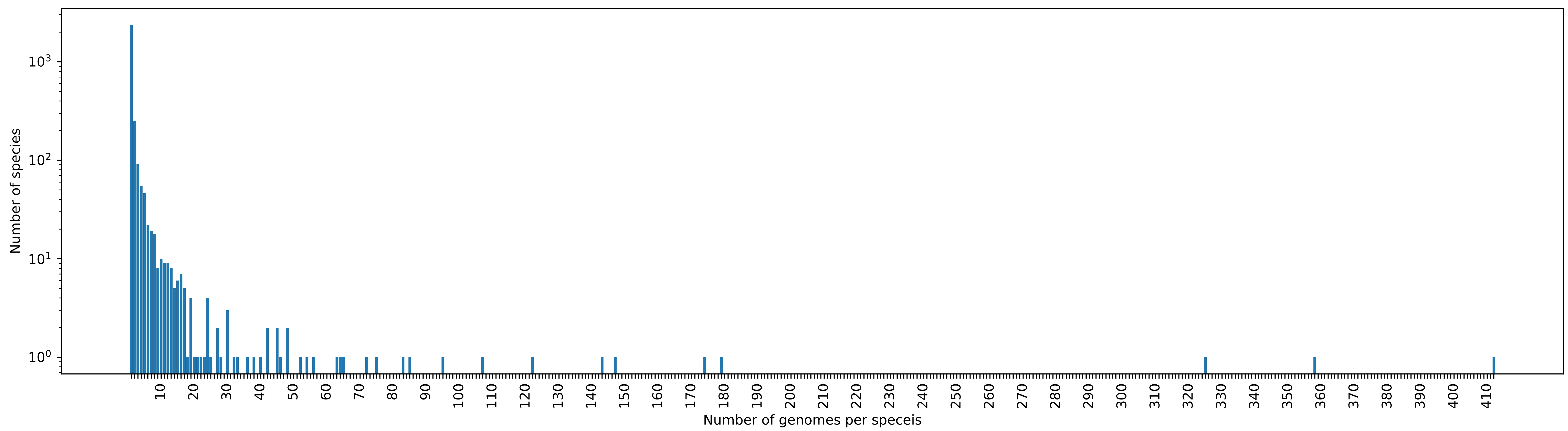

### 2017.pdf

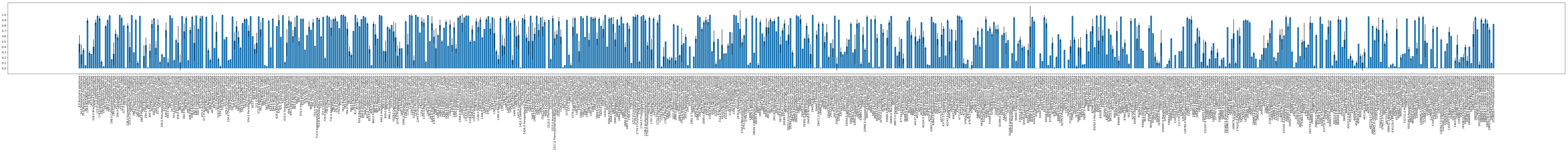

### 2018.pdf

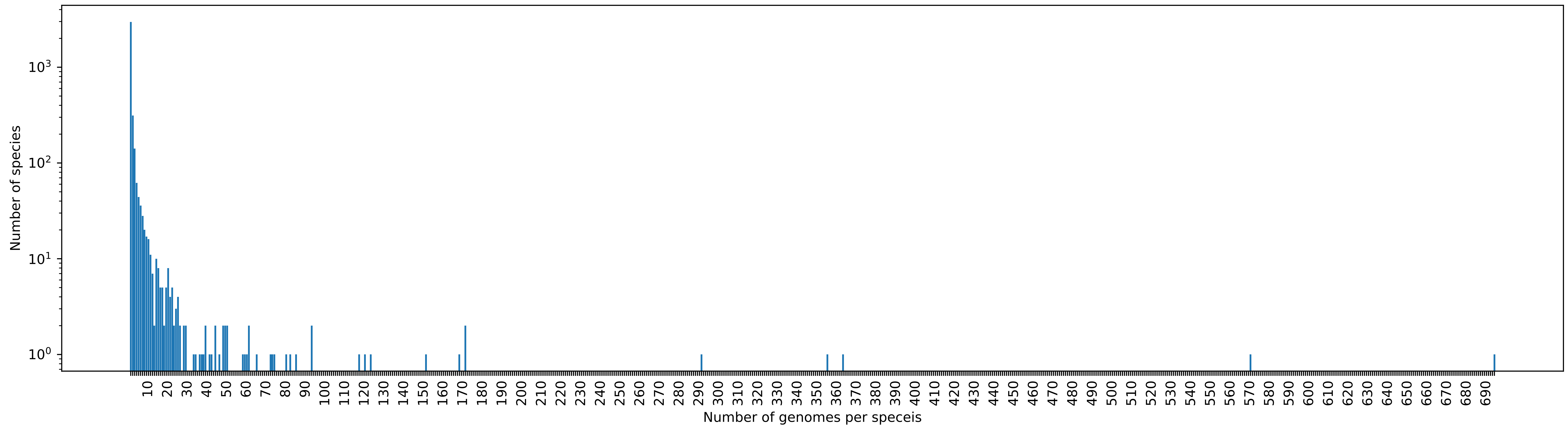

### 2019.pdf

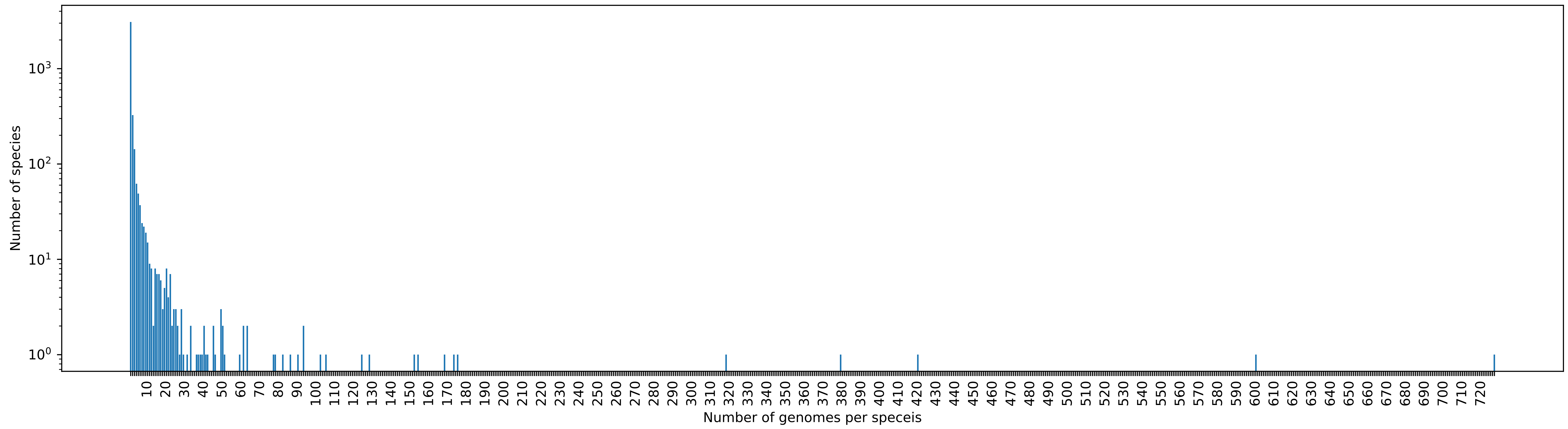

### blast_118110 _against_other_9_no best match.png

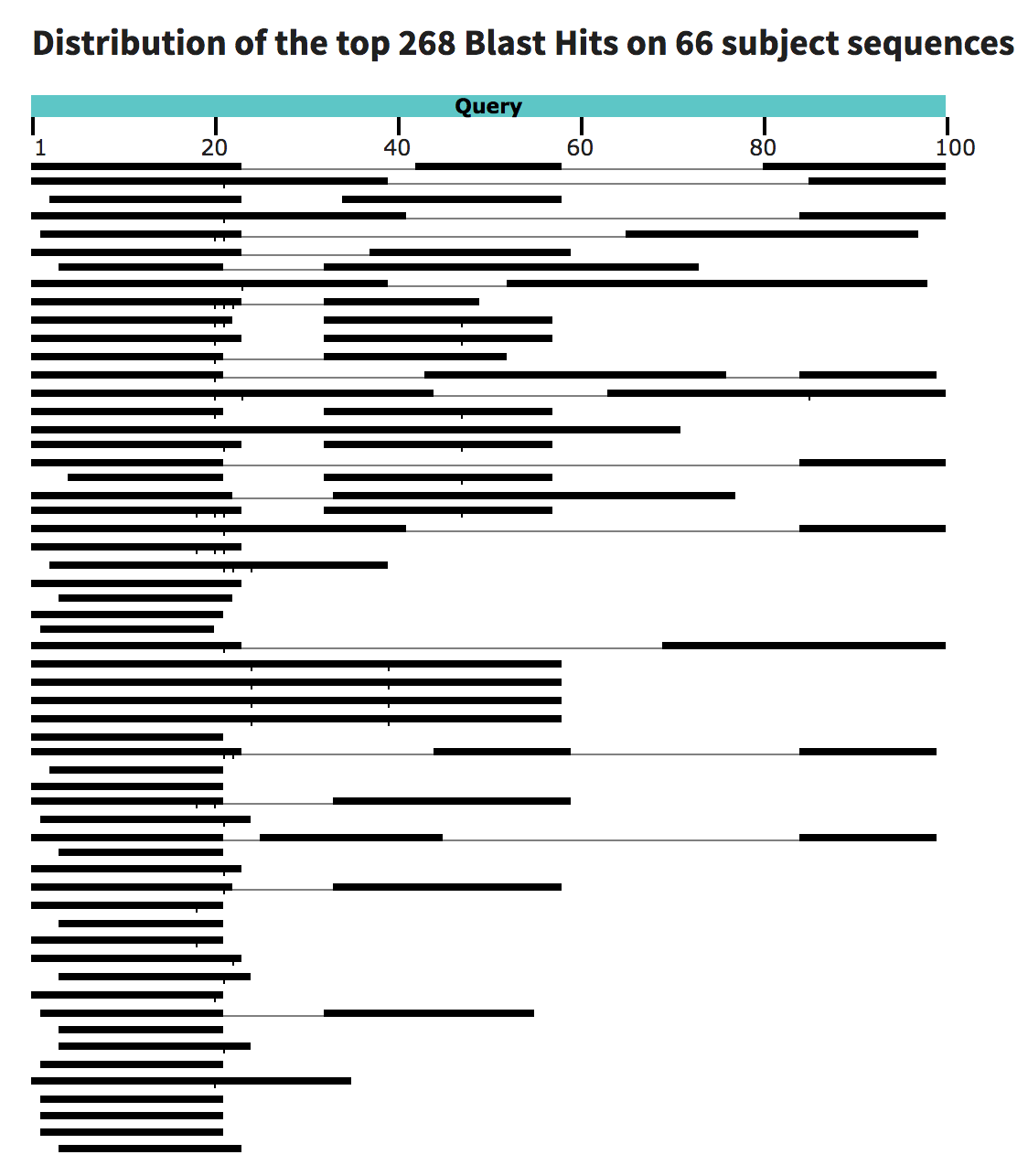

### blast_118110_against_1491.png

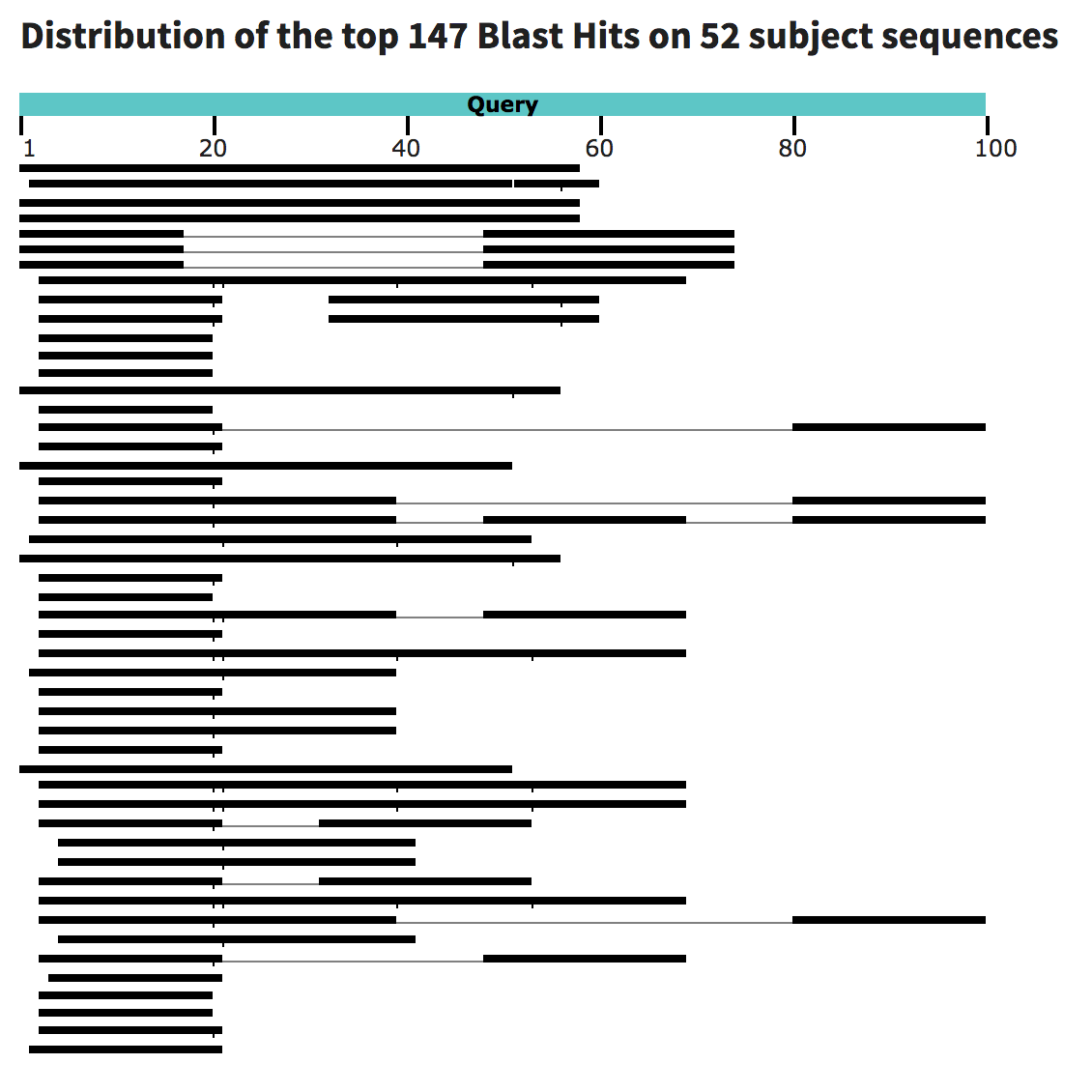

### S4_CLASS_CUMULATIVE_NUM.pdf

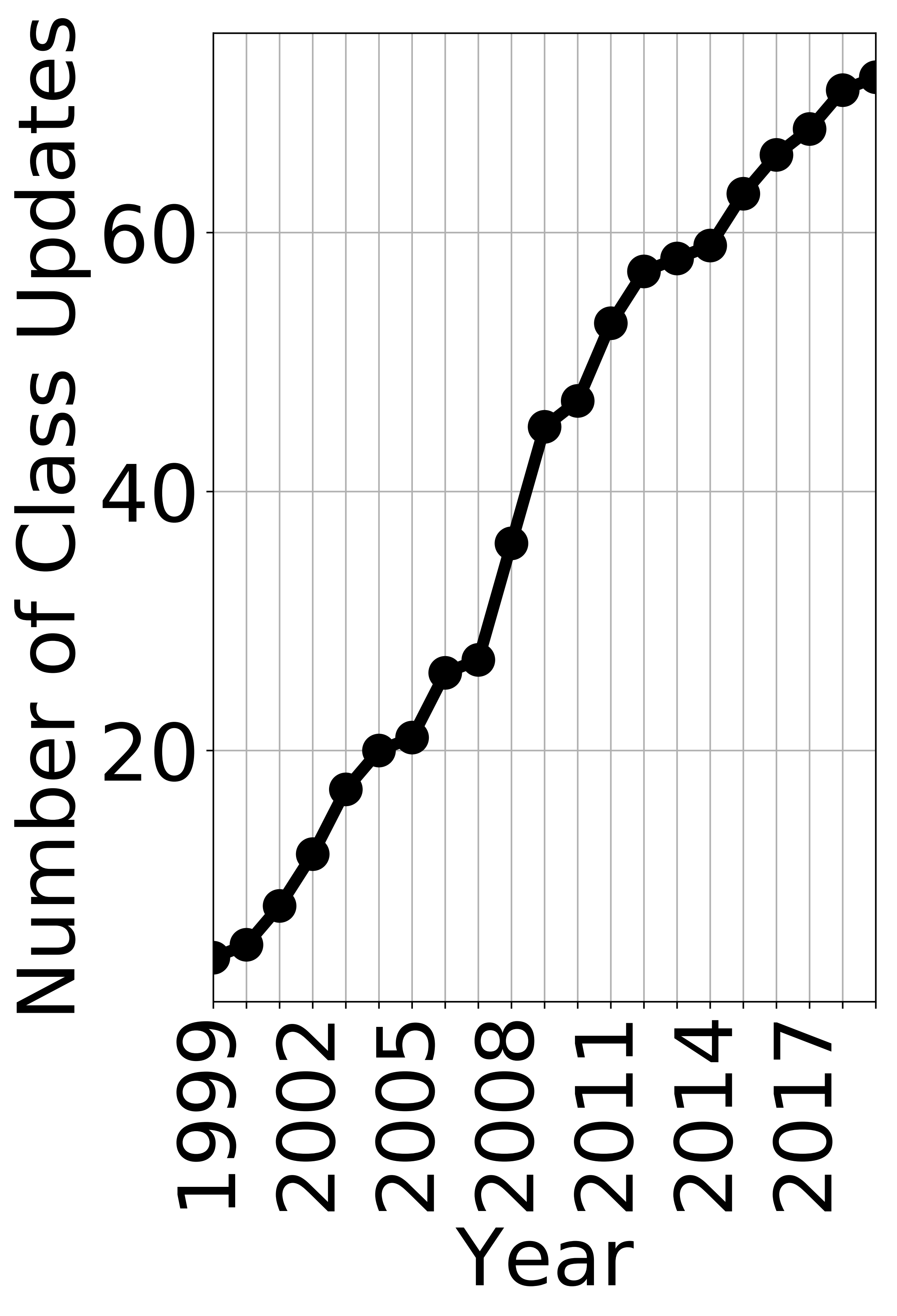

### S4_FAMILY_CUMULATIVE_NUM.pdf

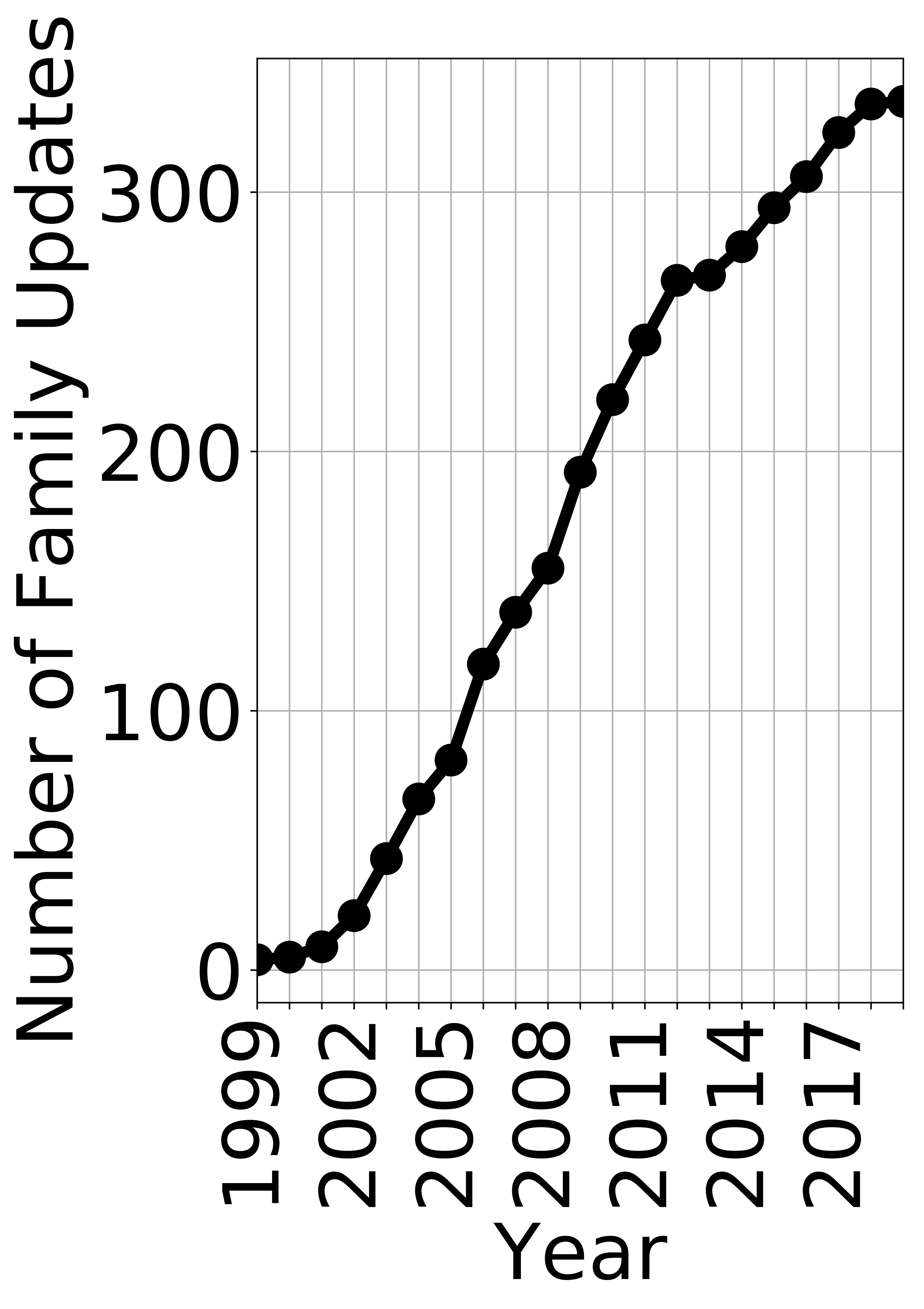

### S4_GENUS_CUMULATIVE_NUM.pdf

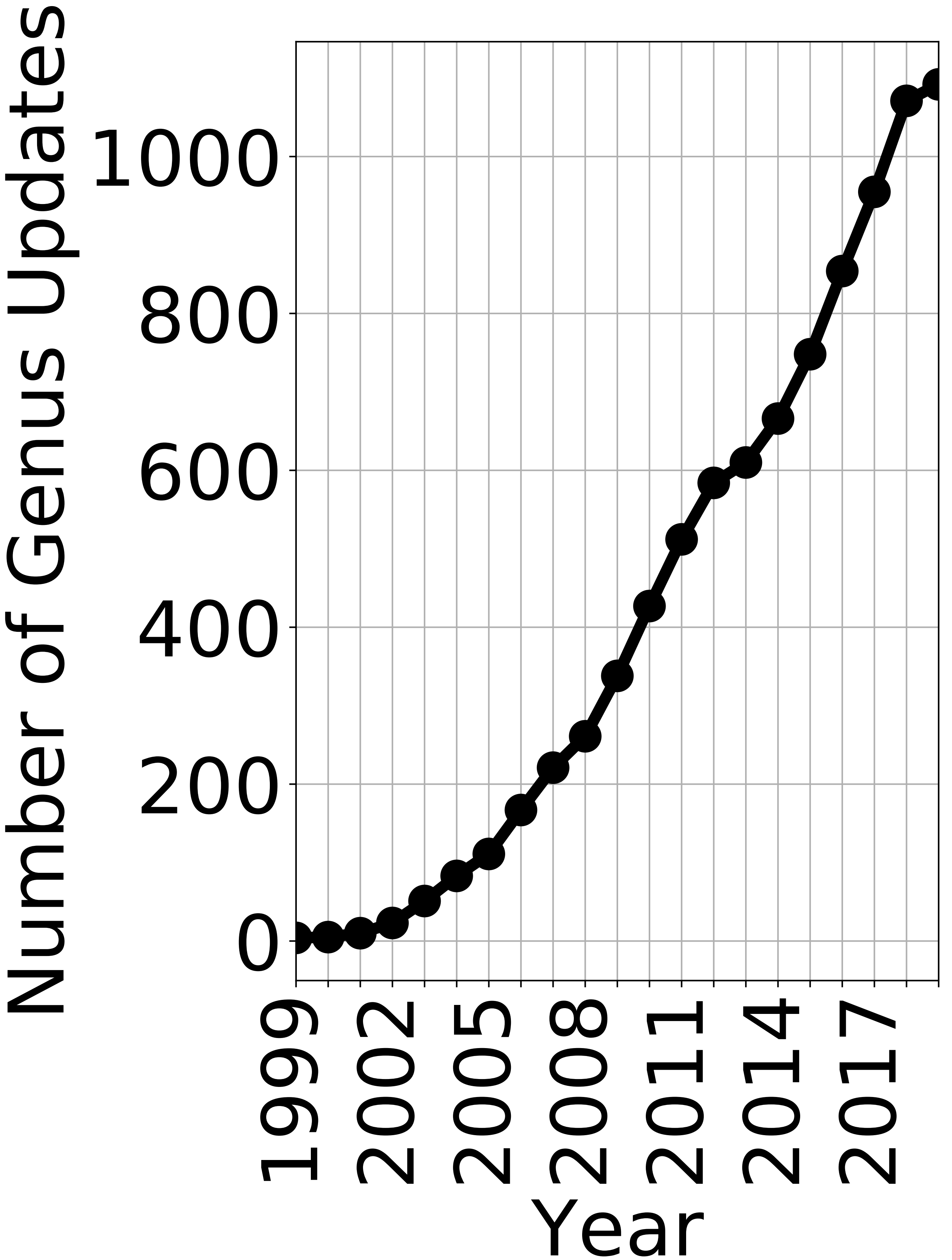

### S4_NEW_UNIQUE_CLASS_UPDATES_NUM.pdf

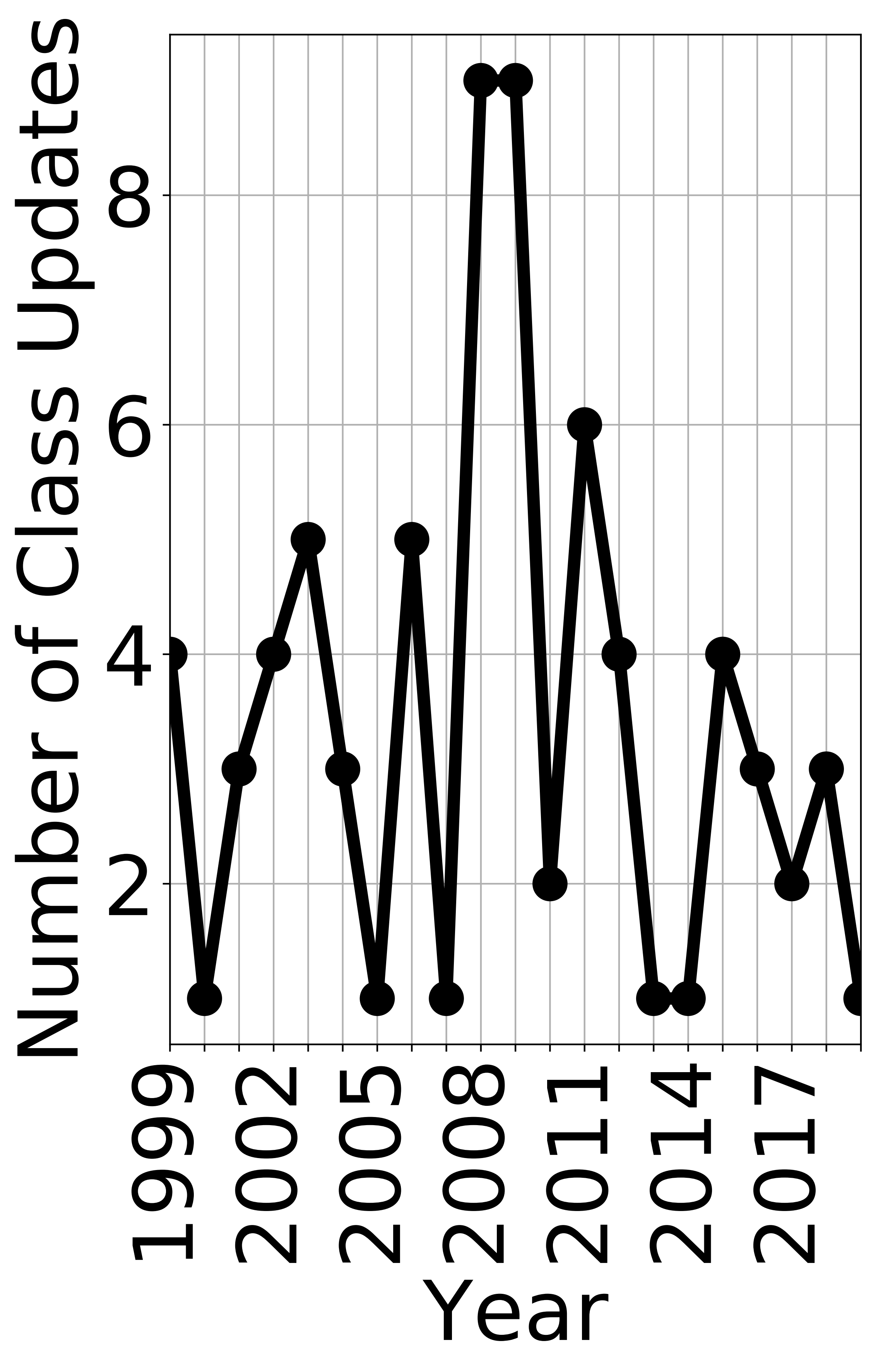

### S4_NEW_UNIQUE_FAMILY_UPDATES_NUM.pdf

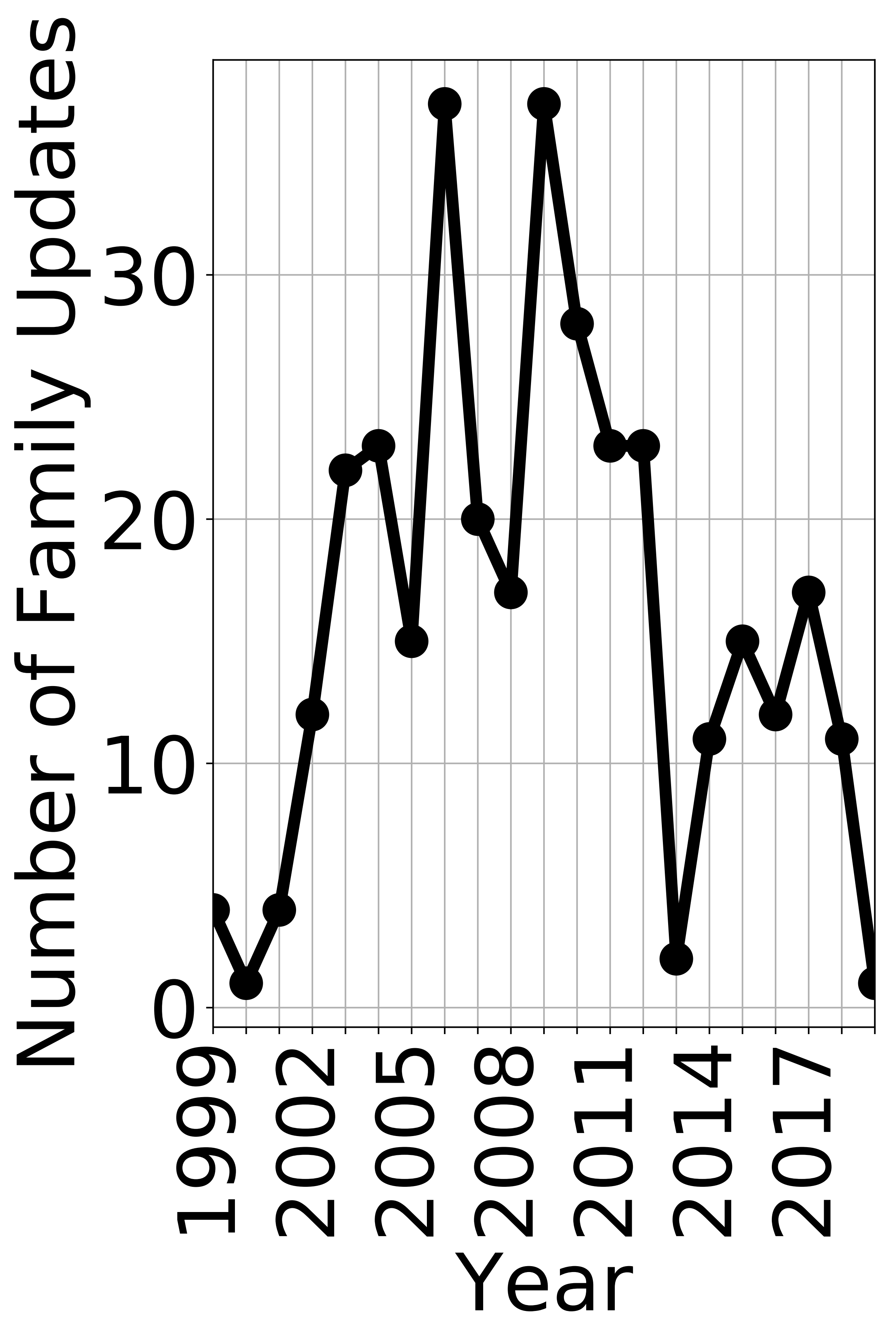

### S4_NEW_UNIQUE_GENUS_UPDATES_NUM.pdf

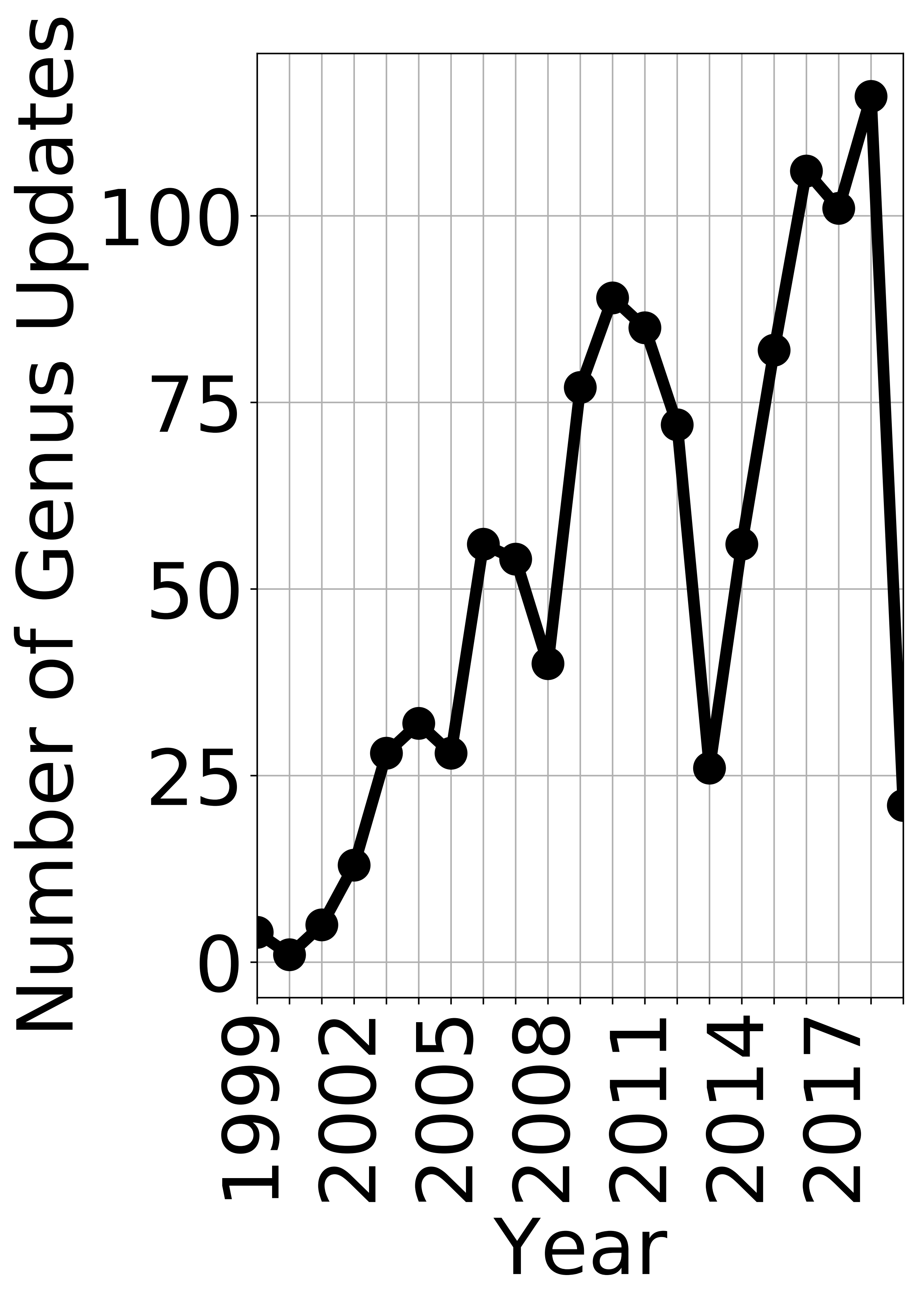

### S4_NEW_UNIQUE_ORDER_UPDATES_NUM.pdf

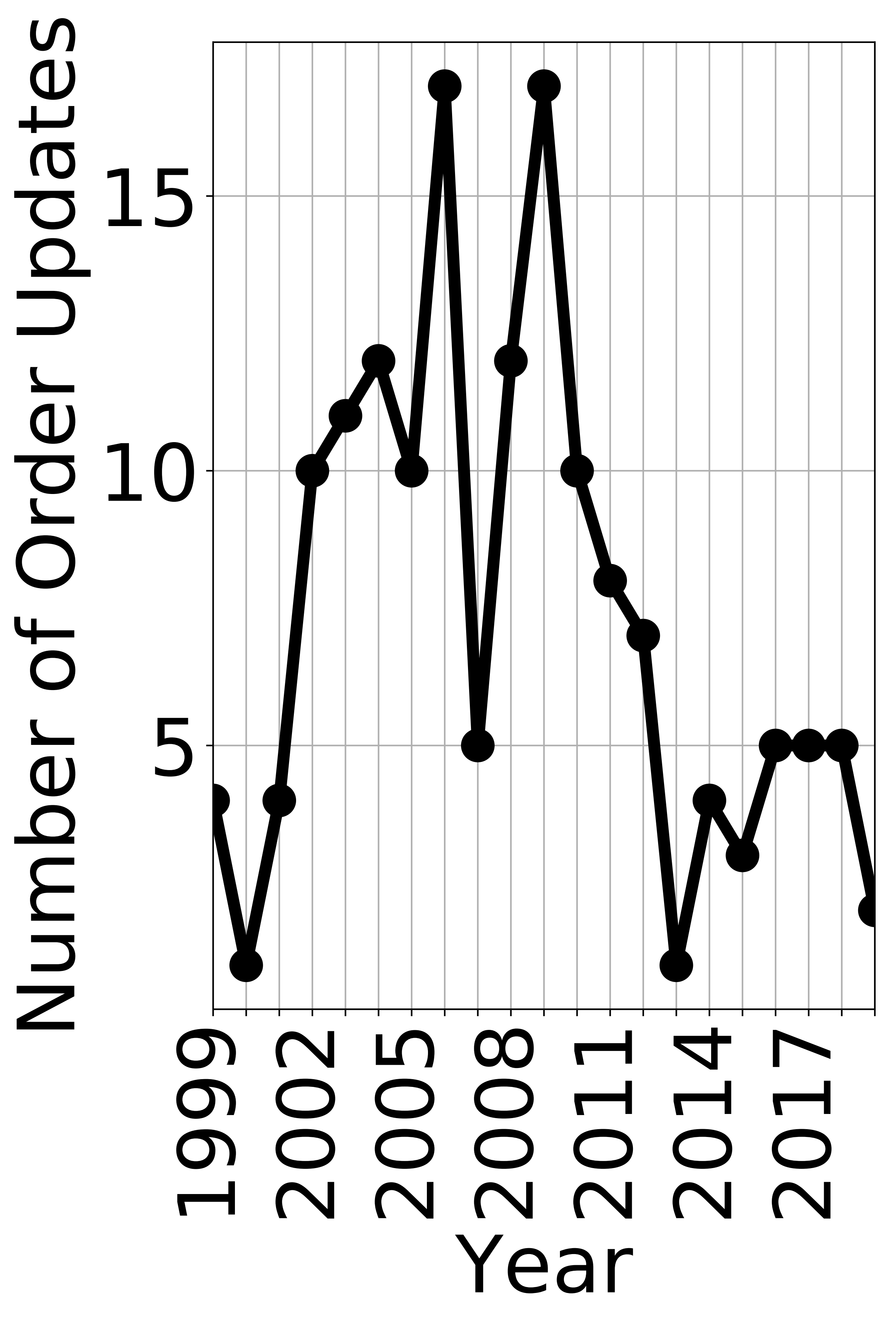

### S4_NEW_UNIQUE_PHYLUM_UPDATES_NUM.pdf

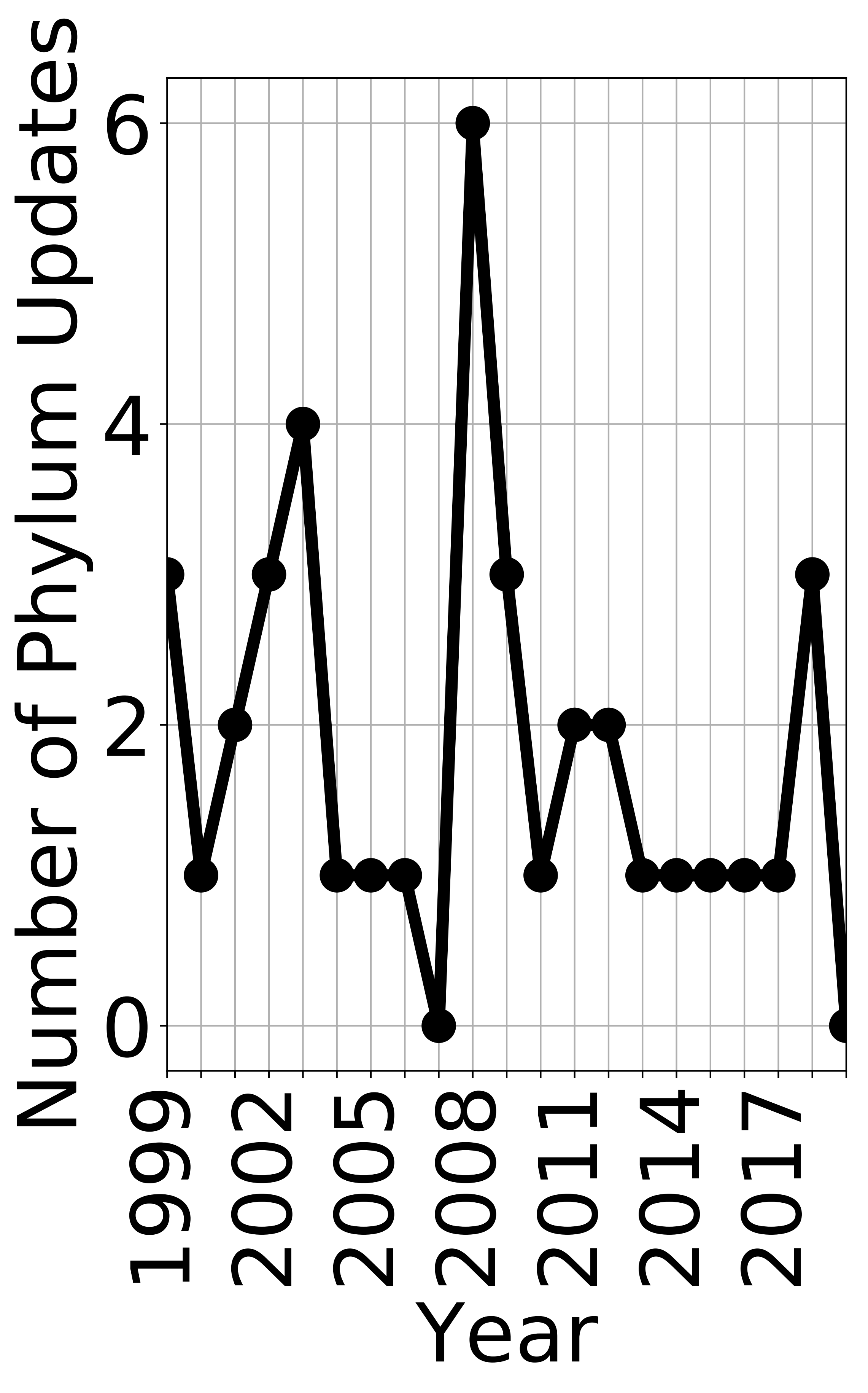

### S4_NEW_UNIQUE_SPECIES_UPDATES_NUM.pdf

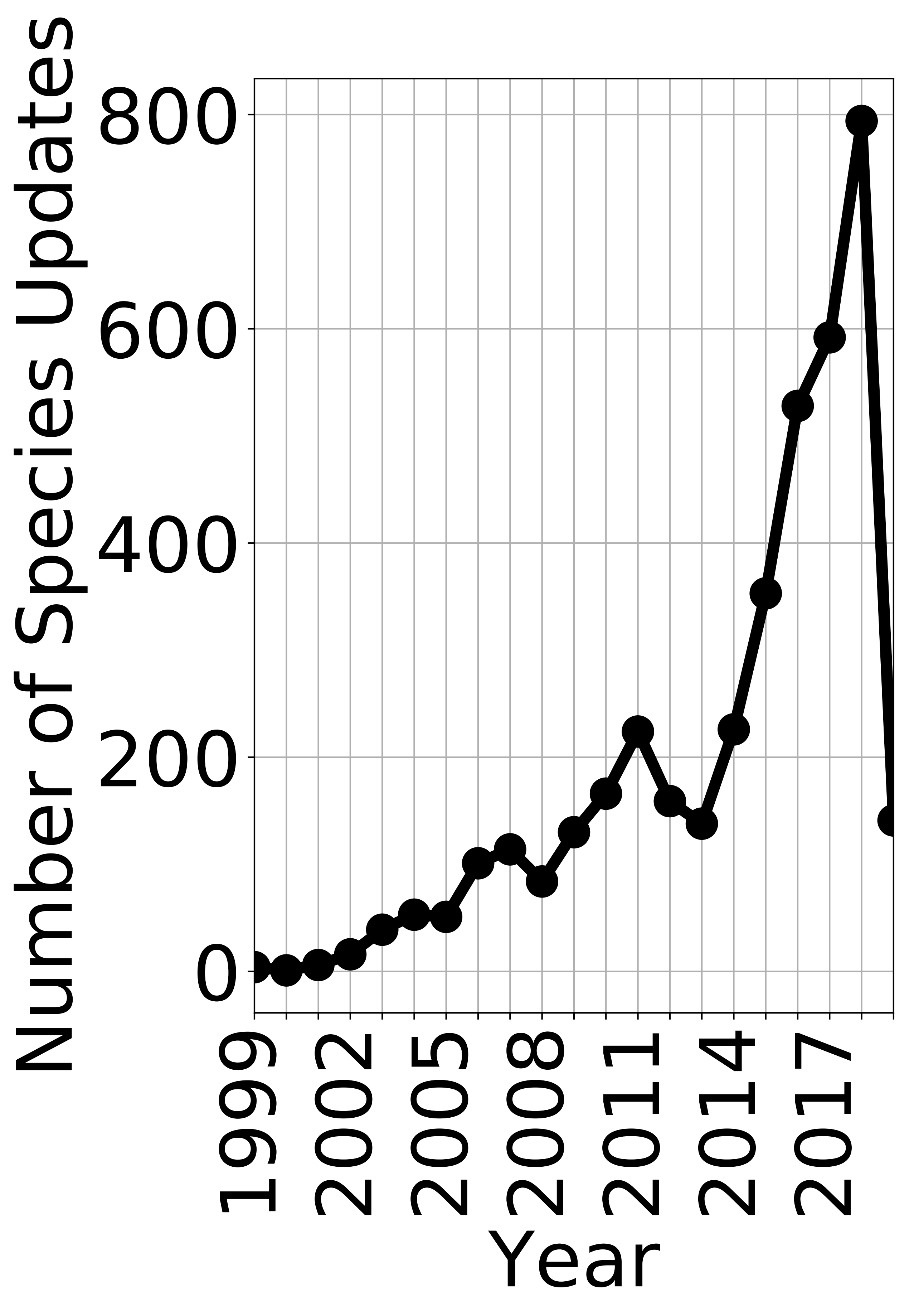

### S4_ORDER_CUMULATIVE_NUM.pdf

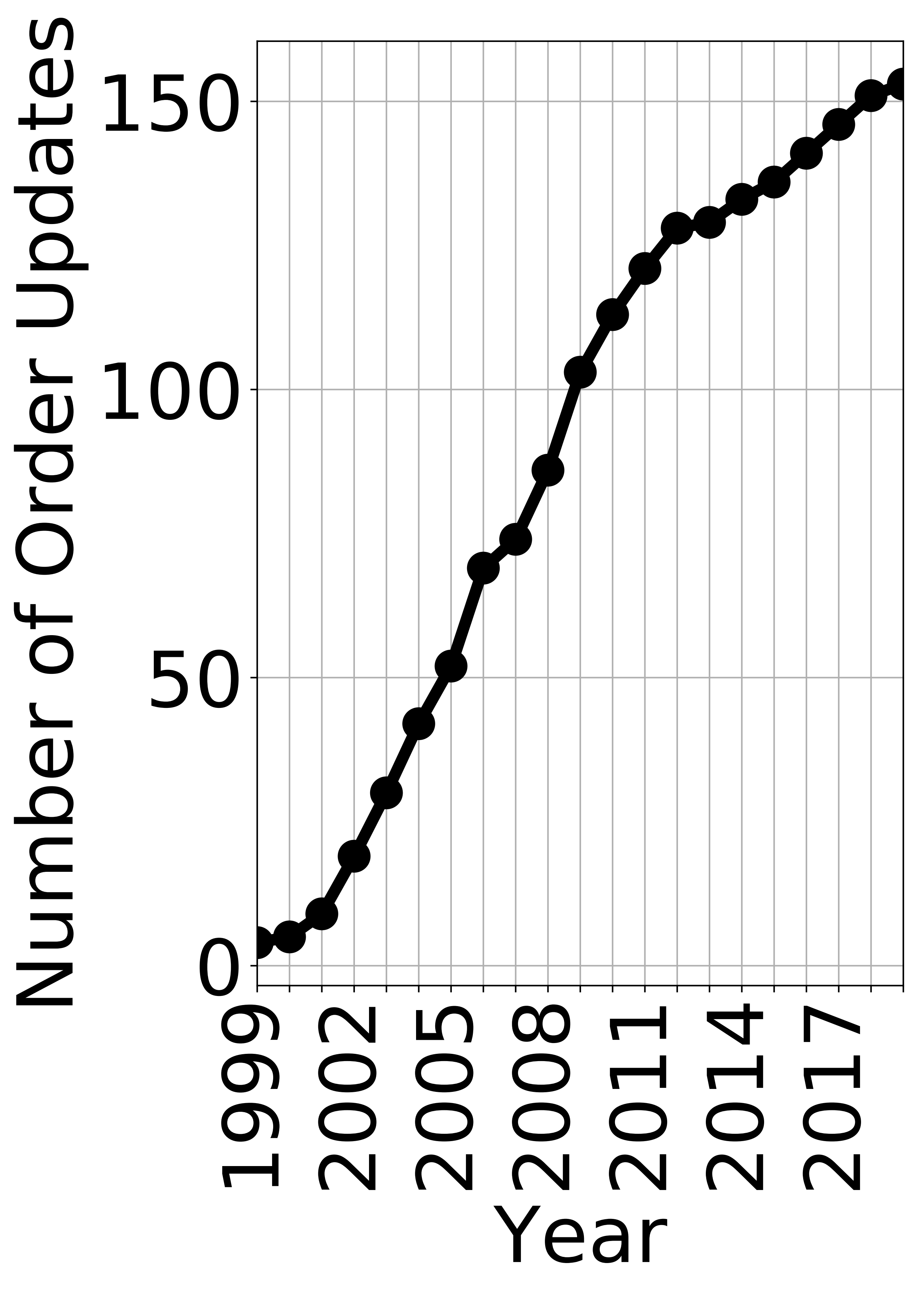

### S4_PHYLUM_CUMULATIVE_NUM.pdf

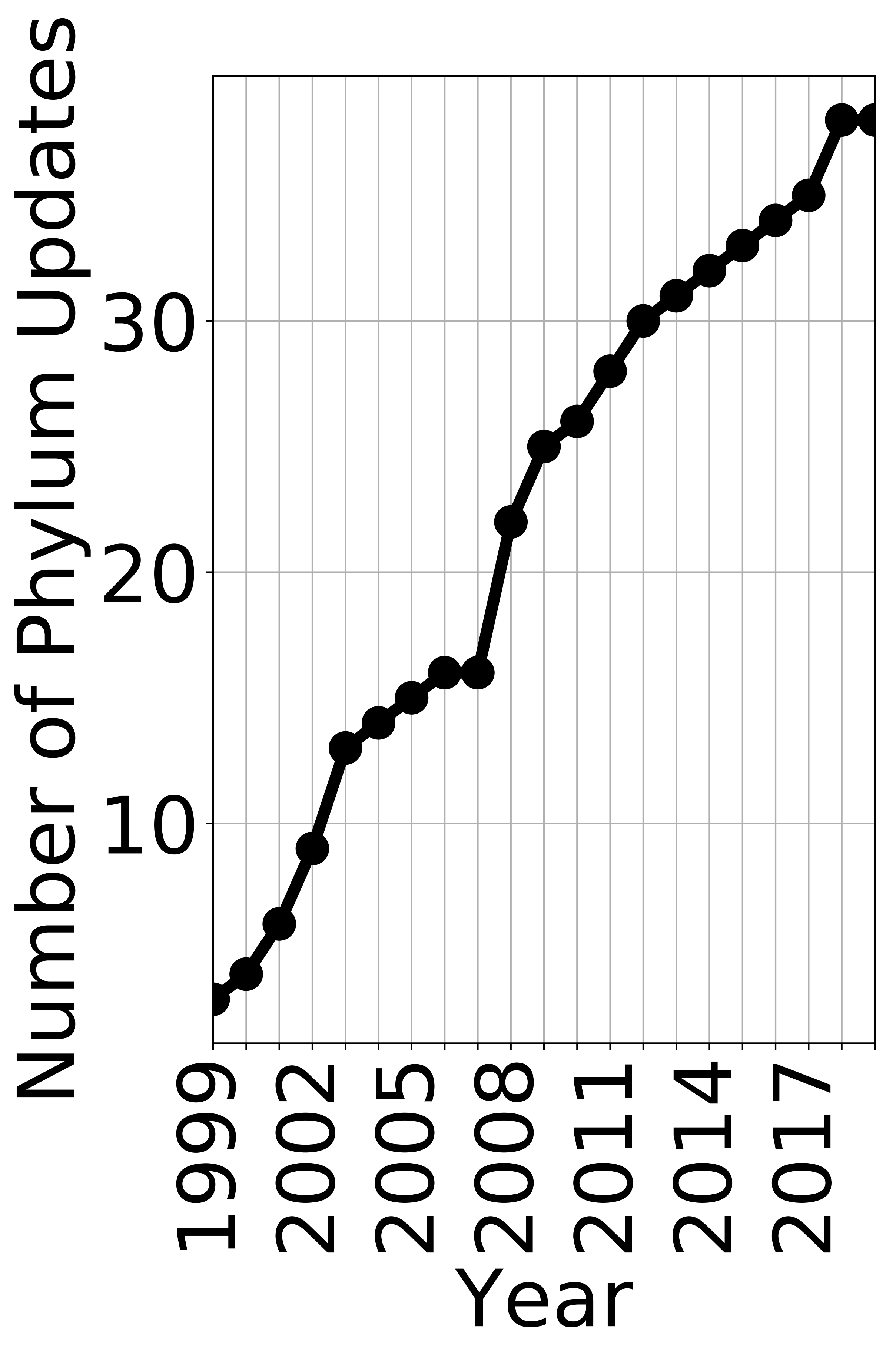

### S4_SPECIES_CUMULATIVE_NUM.pdf

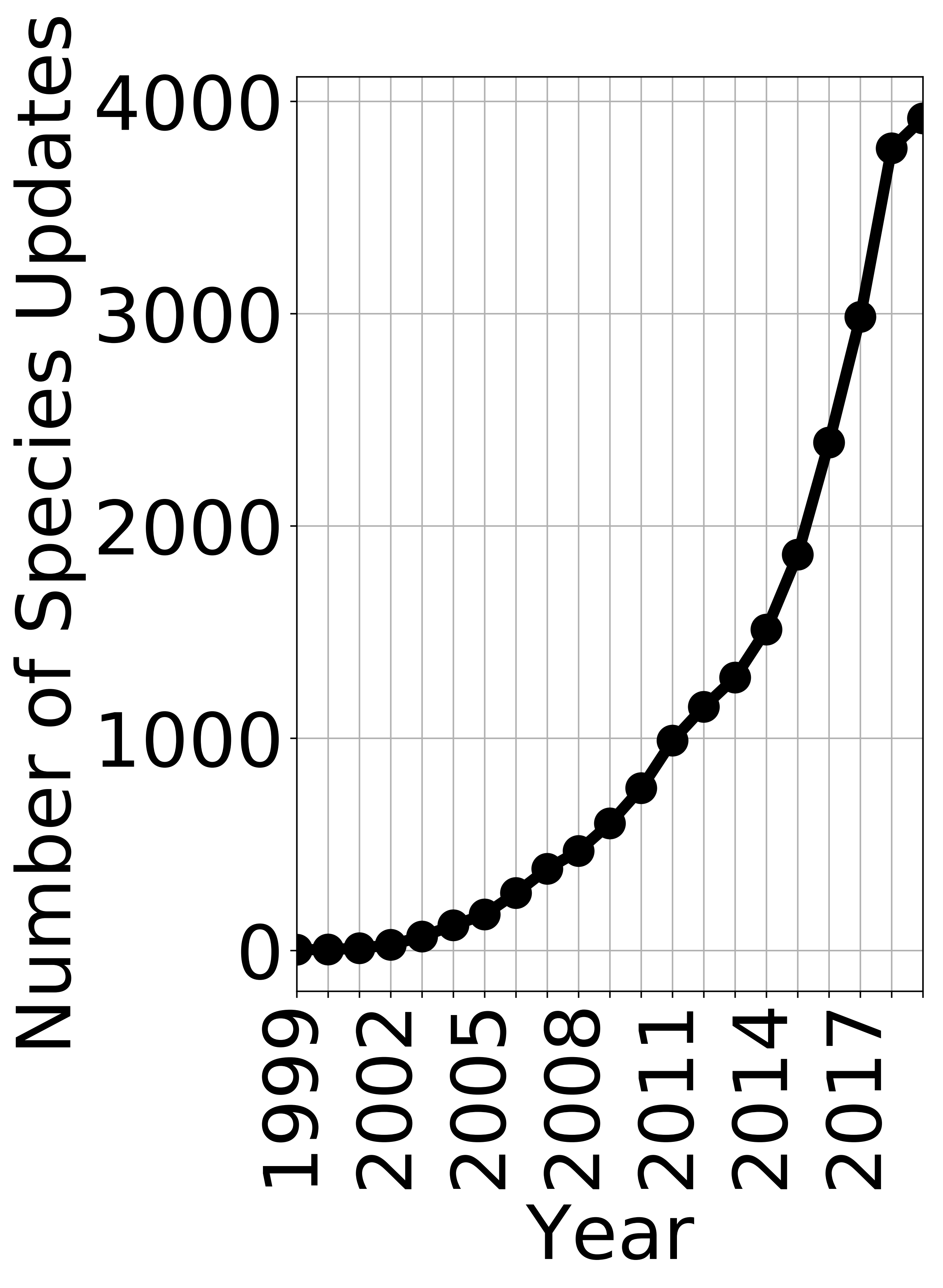

### S7Figure.pdf

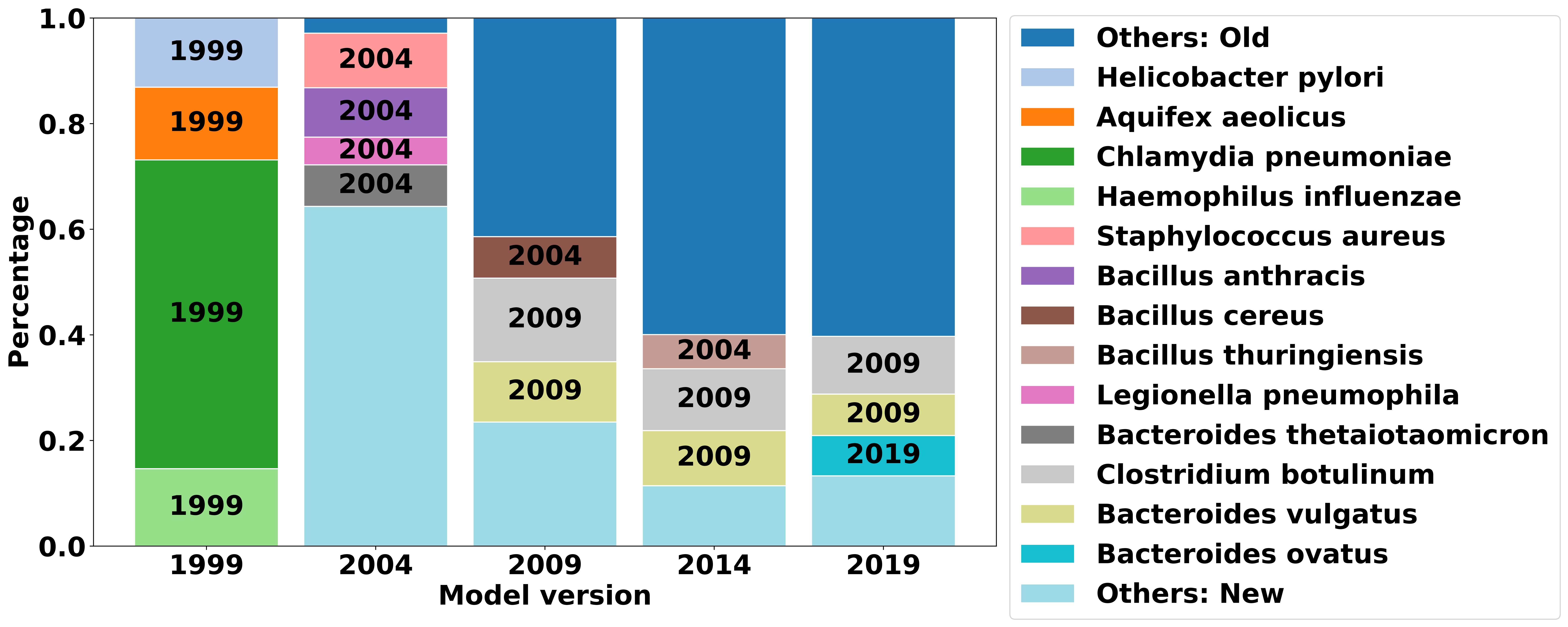

### S8Figure.pdf

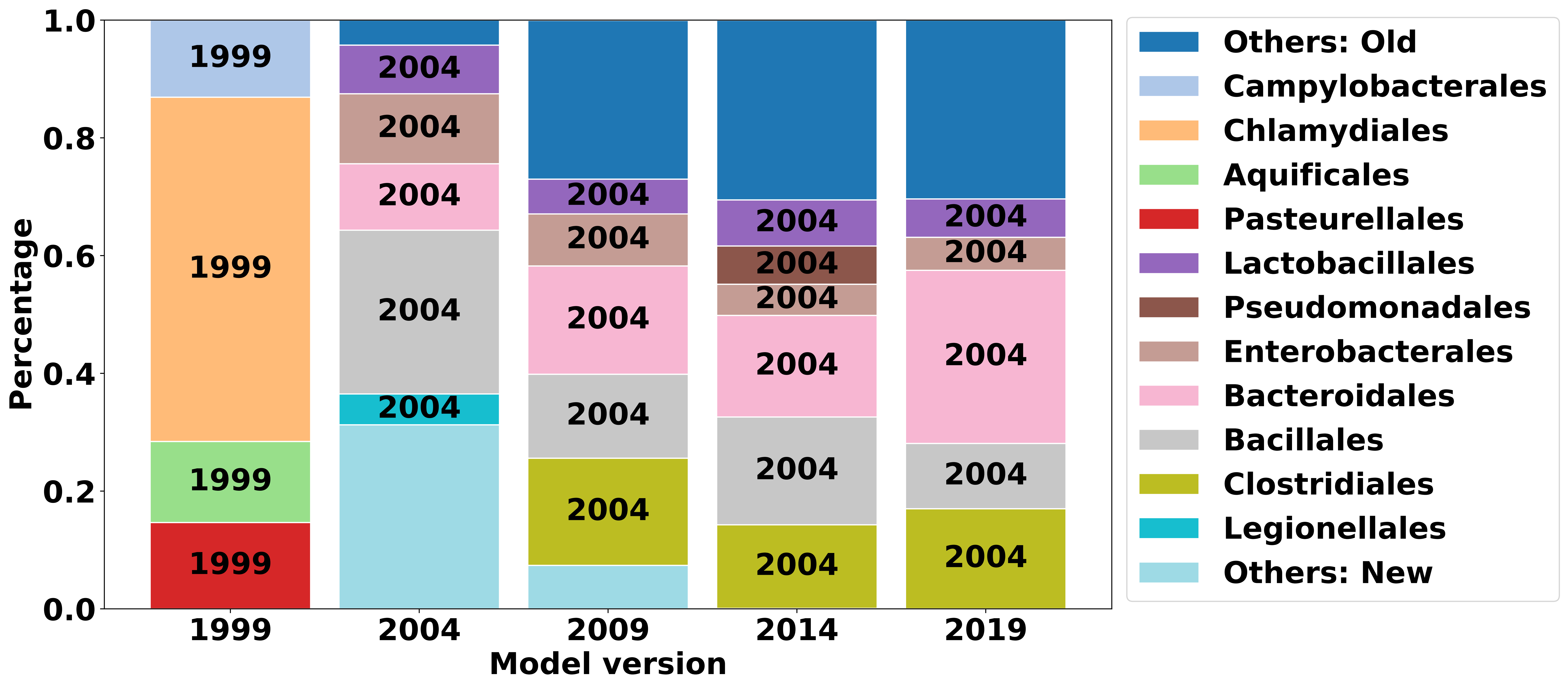
